# SARS-CoV-2 Omicron (BA.1 and BA.2) specific novel CD8+ and CD4+ T cell epitopes targeting spike protein

**DOI:** 10.1101/2022.04.05.487186

**Authors:** Simone Parn, Kush Savsani, Sivanesan Dakshanamurthy

## Abstract

The Omicron (BA.1/B.1.1.529) variant of SARS-CoV-2 harbors an alarming 37 mutations on its spike protein, reducing the efficacy of current COVID-19 vaccines. This study identified CD8+ and CD4+ T cell epitopes from SARS-CoV-2 S protein mutants. To identify the highest quality CD8 and CD4 epitopes from the Omicron variant, we selected epitopes with a high binding affinity towards both MHC I and MHC II molecules and applied other clinical checkpoint predictors including immunogenicity, antigenicity, allergenicity, instability, and toxicity. Subsequently, we found eight Omicron (BA.1/B.1.1.529) specific CD8+ and eleven CD4+ T cell epitopes with a world population coverage of 76.16% and 97.46%, respectively. Additionally, we identified common epitopes across Omicron BA.1 and BA.2 lineages that target mutations critical to SARS-CoV-2 virulence. Further, we identified common epitopes across B.1.1.529 and other circulating SARS-CoV-2 variants, such as B.1.617.2 (Delta). We predicted CD8 epitopes’ binding affinity to murine MHC alleles to test the vaccine candidates in preclinical models. The CD8 epitopes were further validated using our previously developed software tool PCOptim. We then modeled the three-dimensional structures of our top CD8 epitopes to investigate the binding interaction between peptide-MHC and peptide-MHC-TCR complexes. Importantly, our identified epitopes are targeting the mutations on the RNA-binding domain and the fusion sites of S protein. This could potentially eliminate viral infections and form long-term immune responses compared to rather short-lived mRNA vaccines and maximize the efficacy of vaccine candidates against the current pandemic and potential future variants.

## Introduction

The novel Omicron (BA.1/B.1.1.529) variant of SARS-CoV-2, first identified in South Africa in 2021 [**1**], was quickly declared as a variant of concern (VOC) as it had the potential to infect many populations across the world. Although several COVID-19 vaccines and antiviral drugs are currently available, the new variant has challenged public health at an unprecedented scale. Characterized by a heavy mutation load, Omicron has raised alarming immune escape and potentially increased transmissibility compared to the Delta variant [**2**]. B.1.1.529 harbors an alarming number of spike protein substitutions with a total of 15 located on the RNA-binding domain (RBD) of the S protein alone, meaning the new VOC may confer enhanced binding to the human Angiotensin-Converting Enzyme 2 (ACE2) receptor [**2**,**3**]. Moreover, the recent emergence and spread of Omicron sub-lineage BA.2 may pose another risk to public safety. Thus, there is an urgent need for effective booster vaccines against SARS-CoV-2 Omicron variants to reduce the spread of this highly infectious agent.

SARS-CoV-2 spike (S) glycoprotein is a first-generation COVID-19 target due to its structure and function which play an important role in viral infection and pathogenesis [**4**]. Mature S protein consists of two subunits: S1, which includes the RNA-binding domain responsible for the interaction with ACE2, and S2, which facilitates membrane fusion and viral entry [**5**,**6**]. The S1 subunit contains the N-terminal domain (NTD) and the RBD which are the major targets of polyclonal and monoclonal neutralizing antibodies [**7**]. In addition, SARS-CoV-2 hosts a novel protease cleavage site at S1/S2 predicted to increase the virulence of the virus by enabling the cleavage of fusion peptides and therefore promoting entry into lung cells [**4**]. The continued spread of COVID-19 in immunocompromised individuals allows the virus to mutate while generating new SARS-CoV-2 variants. The emergence of B.1.1.529 S protein point mutations and deletions, such as mutations at residues 142-145, 417, 484, and 501, maybe correlated with increased viral transmission and resistance to NTD or RBD antibodies, therefore creating speculation whether the current vaccines provide efficient protection against the virus [**8**,**9**].

The need for a quick and effective vaccine to combat existing and emerging infectious diseases has implemented the use of *in silico* prediction methods, such as immunoinformatics [**10**]. Specifically, immunoinformatics provides a low-cost and rapid approach for epitope-based vaccine design which has many advantages over other conventional types of vaccines. They offer more specific immune responses, promote long-lasting immunity, and reduce undesired side effects [**11**]. Memory T cells specific for SARS-CoV-2 epitopes can persist for 11 years after infection [**12**] compared to relatively short-lived antibody responses [**13**]. Moreover, T cell responses in transgenic mice have been correlated with similar T cell responses in vaccinated and infected humans [**14**,**15**], indicating epitope-based vaccines’ potential to undergo clinical testing at an accelerated rate.

In this study, we obtained immunogenic CD8+ and interferon-gamma (IFNγ) inducing CD4+ T cell epitopes from spike protein of SARS-CoV-2 for a multi-epitope vaccine to protect against the circulating COVID-19 variants and Omicron variant, specifically. Immune Epitope Database (IEDB) [**16**] was utilized to predict both CD8+ and CD4+ T cell epitopes with high binding affinity to MHC class I and II alleles, respectively. Beyond MHC affinity, epitopes were further refined by predicting immunogenicity, population coverage, antigenicity, allergenicity, toxicity, IFNγ secretion, half-life, GRAVY (grand average of hydropathicity), and other amino acid physiochemical properties, to fully maximize the quality and efficiency of the designed vaccine. After predicting CD4+ and CD8+ T cell epitopes, we attempted to identify top immunogenic CD8 peptides (9-10 aa long) within IFNγ inducing CD4 epitopes (15 aa long) that could be used to provoke a strong and long-lasting immune response against the Omicron variant. We validated the predicted epitopes using our previously developed program called PCOptim [**17**]. We then predicted the 3D structures of the final CD8 T cell epitopes and docked them with their respective HLA alleles to visualize the peptide-MHC interaction. Since T-cell receptor (TCR) in complex with pMHC structure is thought to play an important role in the immune response against evading pathogens, we also visualized the pMHC-TCR interaction to gain a better insight into the vaccine mechanism. To safely test the peptide-based vaccine constructs for their efficacy and immunogenicity in pre-clinical trials, we also predicted murine MHC binding affinity to our CD8 peptides.

## Materials and Methods

### Retrieval of SARS-CoV-2 sequence

We retrieved the reference S protein sequence of SARS-CoV-2 Wuhan isolates from the NCBI database using accession number YP_009724390.1. The Omicron BA.1 and BA.2 specific S protein sequences were generated using lineage-defining S protein mutations [**18, 19**].

### CD8+ T cell epitope prediction and immunogenicity modeling

CD8+ T cell epitopes were predicted using NetMHCpan EL 4.1 (IEDB Recommended) [**20-24**]. We predicted CD8+ T cell epitopes for the frequently occurring MHC-I-binding alleles and the HLA allele reference set. The amino acid length of peptides 9.0 and 10.0 was selected as parameters for the identification of MHC I alleles. Subsequently, we selected the top 300 unique epitopes based on rank and predicted their immunogenicity using the IEDB Immunogenicity tool [**25**]. Epitopes with an immunogenicity score >0 were selected for further analysis.

### CD4+ T cell epitope prediction

We used IEDB recommended 2.22 to predict CD4+ T cell epitopes [**26**,**27**]. Full HLA reference set and default epitope length were selected as parameters for the prediction of MHC II binders. We chose the top 300 unique epitopes based on rank for further analysis.

### IFNγ inducing CD4 peptide prediction

Due to the lack of reliable immunogenicity predictor on IEDB.org, the top 300 unique CD4 peptides were refined based on IFNγ inducer properties using the IFNEpitope server [**28**]. IFNEpitope is designed to make predictions based on peptide length, positional conservation of residues, and amino acid composition. Using a hybrid approach of the motif and SVM-based predictions, epitopes returned either positive or negative induction for IFNγ release. Only epitopes that were “positive” for IFNγ release, were selected for further analysis.

### Antigenicity prediction

VaxiJen v2.0 [**29**] was used to predict antigenic CD8 and CD4 epitopes. VaxiJen predicts epitopes based on the physicochemical properties of the protein sequences. A viral model was used for predictions and epitopes that returned a score >0.4 were considered antigenic.

### Allergenicity prediction

AllerTop v2.0 [**30**] was used to predict allergenicity for both CD8 and CD4 epitopes. The server provides an alignment-free method for in silico prediction of allergens and non-allergens based on the physicochemical properties of epitopes. Only “probable non-allergen” peptides were selected for our analysis.

### Toxicity prediction

The toxicity of CD8 and CD4 peptides was determined using the ToxinPred server [**31**]. The method was developed based on machine learning and quantitative matrix using several properties of peptides. The tool returns both toxic and non-toxic peptides. For our analysis, only “non-toxic” epitopes were selected.

### Amino acid physiochemical properties

Evaluation of various physicochemical properties is essential for determining the safety and efficacy of the candidate vaccine. Using ProtParam [**32**] tool on ExPASy, the chemical and physical properties of the top CD8 and CD4 peptides were assessed. Instability index, aliphatic index, GRAVY, and half-life were among the parameters used in the study. The instability index was calculated to determine whether an epitope was stable or unstable in vivo where instability index <40 was selected as the threshold for stable epitopes. Half-life in mammalian reticulocytes in vitro was predicted where half-life >1 hour was selected as a threshold value.

### Worldwide human population coverage analysis

We calculated the population coverage of predicted epitopes with known MHC restriction using the IEDB population coverage tool [**33**]. The percentage of the world population predicted to present our epitopes on their MHC molecules was computed by the tool. The population coverage analysis was performed separately for MHC I and II binders. The analysis covered East Asia, Northeast Asia, South Asia, Southeast Asia, Southwest Asia, Europe, East Africa, West Africa, Central Africa, North Africa, South Africa, West Indies, North America, Central America, South America, and Oceania. Only HLA alleles from the top 300 predicted epitopes ordered according to “rank” were considered in the analysis.

### Murine MHC restriction prediction

We utilized SYFPEITHI [**34**] to predict CD8 peptides’ affinity to murine MHC class I (H2) molecules. The prediction tool is based on published T cell epitopes and MHC ligands and takes into consideration the amino acids in the anchor and auxiliary anchor positions, as well as other frequent amino acids. Two common mouse strains to study COVID-19 vaccines, C57BL/6 and BALB/CJ, were used in mouse MHC affinity predictions. Since scores calculated by the SYFPEITHI server vary, we identified peptides that have been experimentally validated to bind to C57BL/6 and BALB/CJ supertypes. Subsequently, we compared the scores of reference peptides: RTFSFQLI [**35**], MYIFPVHWQF [**36**], and RPQASGVYM [**37**] to our predicted CD8 epitopes.

### Validation using our Population Coverage Optimization Software

We earlier developed a program called PCOptim to validate the predicted epitopes. The program can be used to generate an optimized dataset of epitopes with maximum population coverage [**17**]. The tool accepts the input of several epitope-HLA allele pairs and generates an optimized list of epitopes that provide the maximum possible population coverage. PCOptim produces the optimized list of epitopes by conserving all unique HLA alleles, thereby ensuring that population coverage remains maximized. However, the program does not consider several clinical checkpoint variables: antigenicity, allergenicity, or toxicity. The set of epitopes that pass all clinical checkpoint parameters will be more limited and will likely have lower population coverage than the optimized dataset. Therefore, the optimized set of epitopes can be used as a point of comparison to identify geographic regions that may receive lower coverage.

### Three-dimensional (3D) structure prediction

Three-dimensional (3D) structure prediction plays a useful role in illustrating the binding of the peptide to a respective MHC molecule. To visualize the peptide-MHC complex, we first predicted the tertiary structure of the top CD8 epitopes using PEPstrMOD [**38**]. The 3D structure of the HLA allele that the S protein epitope binds to was downloaded from Protein Data Bank on <u>RCSB.org</u>. The native peptides from the RCSB structures were replaced with our predicted epitopes using PyMol. We then spatially docked the predicted epitope in the MHC groove using PyMOL and FlexPepDock [**39**]. The best model provided by the server was energy minimized using HADDOCK 2.4 [**40**]. We also modeled the 3D structure of the peptide-MHC-TCR complex using TCRmodel [**41**].

## Results

### Workflow of T cell predictions

A schematic presentation of the immunoinformatic methods was proposed for epitope predictions as described in our previous study (**Figure 1**) [**42**]. We started the analysis by creating SARS-CoV-2 Omicron (BA.1 and BA.2) specific S protein sequences where lineage-defining mutations were replaced in the SARS-CoV-2 Wuhan reference sequence. We used experimentally validated prediction tools from our previous study [**43**] to predict CD8 and CD4 epitopes with MHC Class I and II binding affinity, respectively. We applied additional parameters to obtain the highest quality epitopes for the multi-epitope vaccine. World population coverage was thereafter calculated to assess the protection scope of selected epitopes. To test the efficacy of the epitopes in preclinical trials, we predicted the binding affinity of our CD8 epitopes to murine MHC alleles. To validate the CD8 epitopes, we utilized our earlier developed program called PCOptim. Eventually, we visualized peptide-MHC (pMHC) and pMHC-TCR interactions in three-dimensional models to give a better insight into T cell-mediated immune response mechanism.

**Figure 1.**
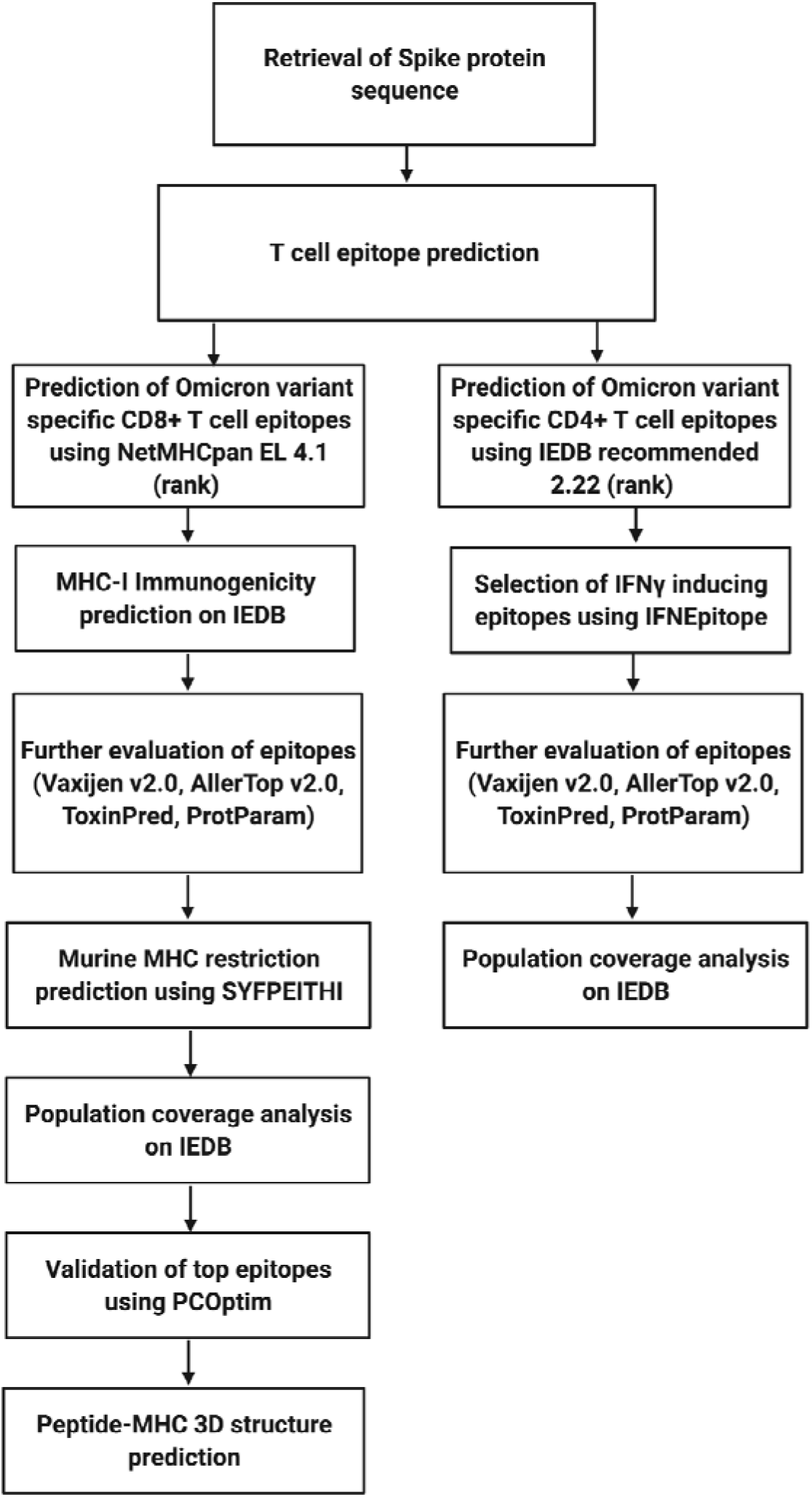
Flowchart of the T cell epitope prediction mechanism. Omicron variant-specific CD8+ and CD4+ T cell epitopes were predicted using freely available prediction tools on Immune Epitope Database (IEDB). MHCI binders were refined by immunogenicity and MHCII binders by IFNγ inducer capability. All CD8 and CD4 epitopes were then evaluated and selected based on their antigenicity, allergenicity, toxicity, and physicochemical properties. Population coverage was computed for both CD8 and CD4 epitopes with known MHC restriction. CD8 peptides were further validated using our software PCOptim. Murine MHC binding affinity to top CD8 epitopes was predicted and the peptide-MHC structures were modeled in three-dimensional analysis.

### SARS-CoV-2 Omicron variant mutations

The Omicron variant, also known as PANGO lineage BA.1 or B.1.1.529, harbors up to 59 mutations in the entire genome with 37 of them located on the S protein, the mediator of host cell entry (**Figure 2**). Among these, Omicron hosts a novel 3 amino-acid insertion at position 214. The RBD region which is considered the main target of neutralizing antibodies as it binds the human ACE2 receptor contains 15 of these mutations. Besides novel mutations in the Omicron BA.1 variant, some of the mutations are also observed in other SARS-CoV-2 variants. The Omicron sub-lineages BA.1 and BA.2 differ in some of the mutations, including the spike protein, however, the two have at least 20 common mutations (**Figure 2**). B.1.617.2 variant shares 3 amino acid mutations with B.1.1.529: G142D, T478K, and D614G whereas B.1.351 and P.1 share mutations in residues K417N, E484K, and N501Y which are correlated with increased binding affinity to ACE2 [**44, 45**].

**Figure 2.**
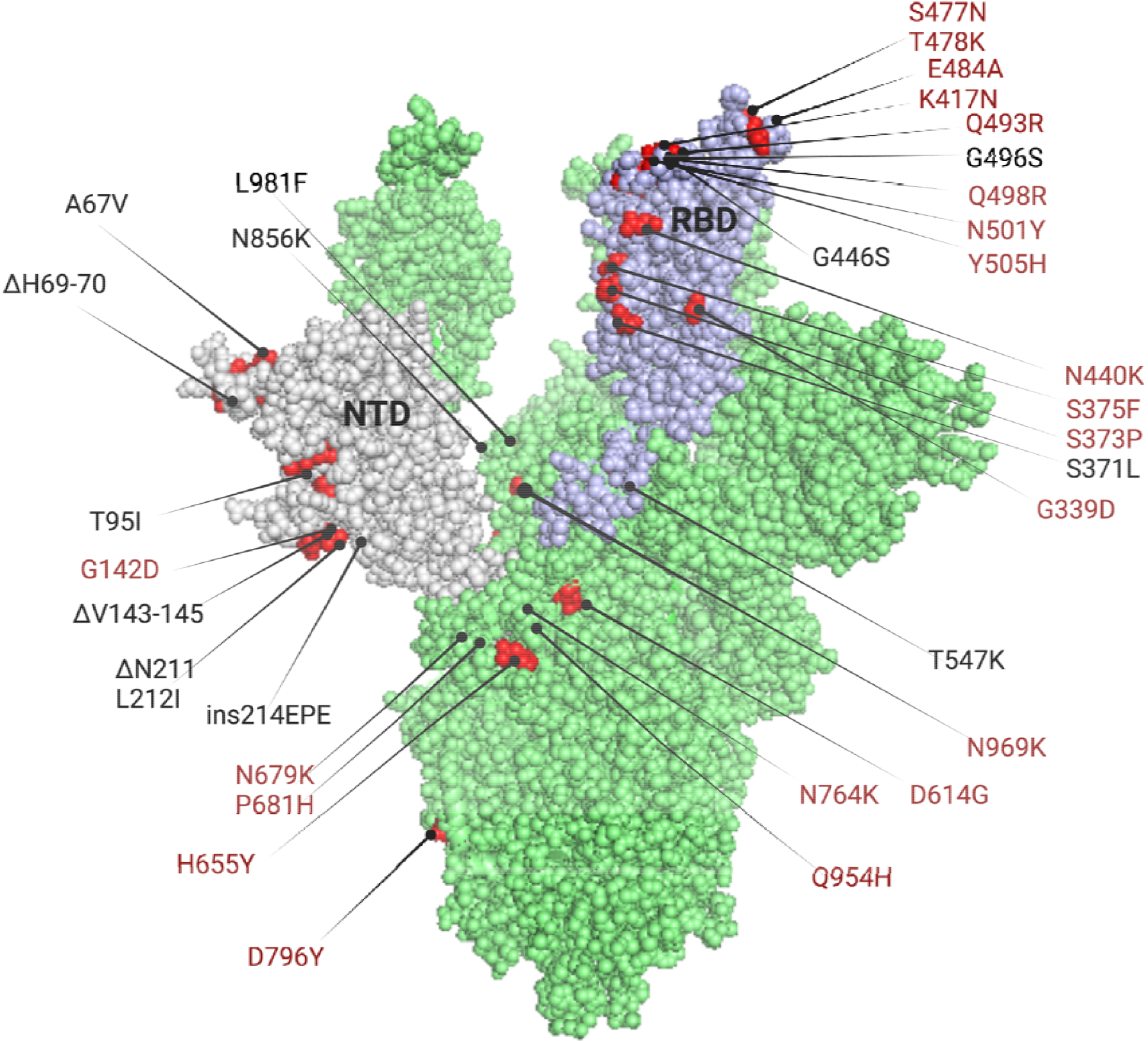
3D structure of SARS-CoV-2 S protein (PDB: 7VNE) highlighting the mutational landscape of Omicron variant (BA.1/ B.1.1.529). Omicron variant harbors 37 mutations on the S protein with most of the mutations located on the RNA-binding domain. Mutations highlighted in red are shared between BA.1 and BA.2 sub-lineages.

### CD8+ T cell epitope prediction

We used NetMHCpan EL 4.1 based on rank, as well as additional parameters including MHC-I immunogenicity predictor, VaxiJen 2.0, AllerTop 2.0, ToxinPred, and ProtParam, to select only immunogenic, antigenic, non-allergenic, non-toxic, and stable CD8+ T cell epitopes from SARS-CoV-2 Omicron (BA.1) S protein. The search yielded 25 immunogenic, antigenic, non-allergenic, non-toxic, and stable epitopes. We then compared these epitopes to other SARS-CoV-2 variant CD8 peptides, identified by our previous study [**43**]. This led us to identify 12 of 25 common epitopes among Omicron (BA.1/B.1.1.529), Alpha (B.1.1.7), Beta (B.1.351), Gamma (P.1), Delta (B.1.617.2), US variants (S protein mutations), and Cluster 5 mink variants (**Table 1, Supplementary Table S1**) which could be used in a multi-epitope vaccine to target multiple variants simultaneously. The remaining 13 CD8 peptides appeared to be matching with the S protein reference sequence of SARS-CoV-2 indicating the absence of any BA.1 variant-specific epitopes.

**Table 1.**
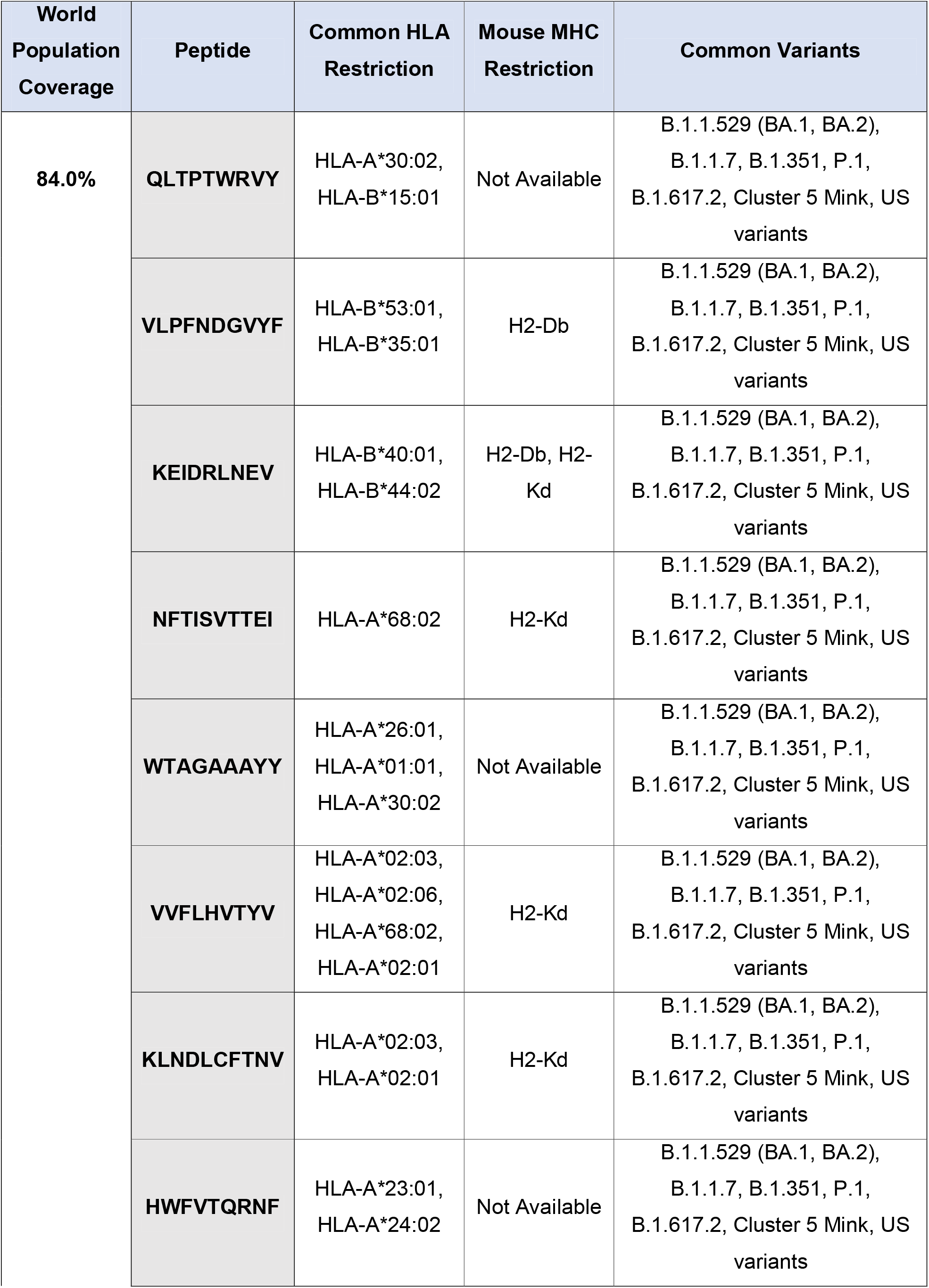
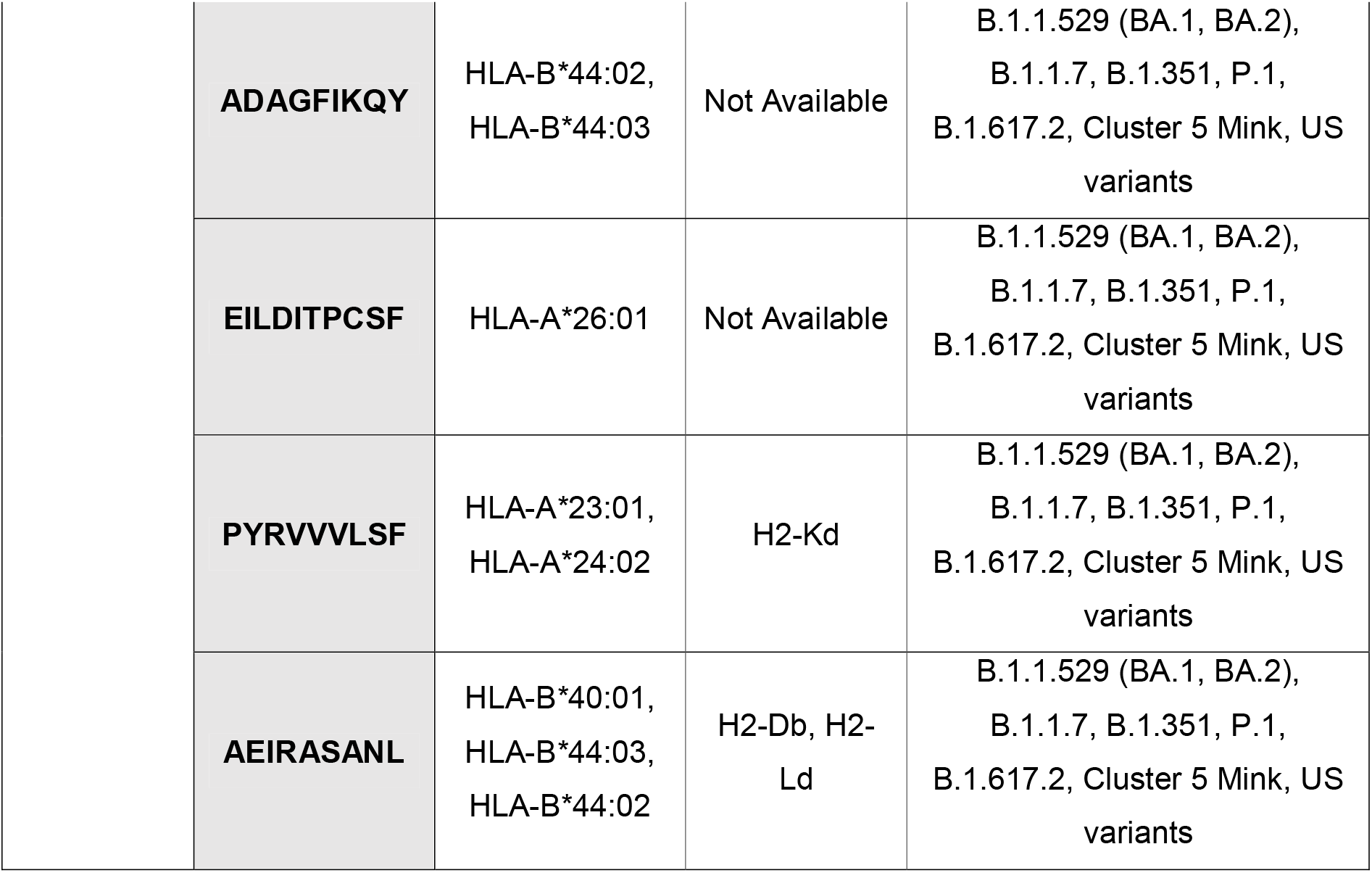
Common S protein CD8+ T cell epitopes among Omicron sub-lineages BA.1 and BA.2 as well as other SARS-CoV-2 variants including Alpha (B.1.1.7), Beta (B.1.351), Gamma (P.1), Delta (B.1.617.2), US variants (S protein mutations), and Cluster 5 mink variants. All predicted epitopes are immunogenic, antigenic, non-allergenic, non-toxic, and stable. The world population coverage of the selected epitopes is also presented.

The stringent method provides an efficient way to match epitopes with 100% sequence similarity. We compared all predicted immunogenic reference epitopes to immunogenic BA.1 epitopes. Successively, the search returned 25 BA.1 specific immunogenic epitopes where 8 were identified to be immunogenic, antigenic, non-allergenic, non-toxic but unstable in vivo (**Table 2, Supplementary Table S2**). We also predicted CD8 peptides specific to the stealth version of the Omicron variant of the BA.2 sub-lineage. The study yielded three immunogenic antigenic non-allergenic non-toxic and stable epitopes specific to BA.2 sub-lineage (**Supplementary Table S3**). Using our 8 BA.1 specific CD8 epitopes, we aimed to identify any common epitopes among BA.1 and BA.2 sub-lineages. The search yielded one CD8+ T cell epitope (KSHRRARSV) present in both Omicron lineages and located at the furin cleavage site. Finally, we checked if any BA.2 CD8 epitopes were also common to other SARS-CoV-2 strains. Successively, BA.2 shares identical epitopes with all other SARS-CoV-2 variants listed in **Table 1**.

**Table 2.** Omicron (BA.1/B.1.1.529) specific S protein CD8+ T cell epitopes and their world population coverage. All predicted epitopes are immunogenic, antigenic, non-allergenic, and non-toxic. Variant-specific mutations are written in red color. We identified one common CD8+ T cell epitope (KSHRRARSV) across Omicron BA.1 and BA.2 sub-lineages.

| World Population Coverage | Peptide | Targeted S Mutations | HLA Restriction | Mouse MHC Restriction | Omicron sub-lineage |
| --- | --- | --- | --- | --- | --- |
| 76.16% | N**LAPFF**TFK | S371L,<br>S373P,<br>S375F | HLA-A*11:01,<br>HLA-A*03:01,<br>HLA-A*68:01 | Not Available | BA.1 |
|  | Y**NLAPFF**TF | S371L,<br>S373P,<br>S375F | HLA-A*23:01,<br>HLA-A*24:02 | H2-Ld, H2-Db | BA.1 |
|  | G**VYFASIE**K | T95I | HLA-A*11:01,<br>HLA-A*03:01,<br>HLA-A*30:01 | Not Available | BA.1 |
|  | **KSHRR**ARSV | N679K,<br>P681H | HLA-A*30:01 | H2-Kd | **BA.1, BA.2** |
|  | **APFF**TFKCY | S373P,<br>S375F | HLA-B*35:01 | H2-Ld | BA.1 |
|  | V**LYNLAPFF** | S371L,<br>S373P,<br>S375F | HLA-A*32:01,<br>HLA-A*23:01 | Not Available | BA.1 |
|  | **REPE**DLPQGF | ins214EPE | HLA-B*44:03,<br>HLA-B*44:02 | H2-Db | BA.1 |
|  | **RSYSFR**PTY | Q493R,<br>G496S,<br>Q498R,<br>N501Y | HLA-A*30:02,<br>HLA-A*32:01,<br>HLA-A*30:01,<br>HLA-B*57:01,<br>HLA-B*15:01,<br>HLA-B*58:01,<br>HLA-A*03:01,<br>HLA-A*11:01 | Not Available | BA.1 |

### CD4+ T cell epitope prediction

Using IEDB recommended 2.22 based on rank, IFNEpitope, VaxiJen 2.0, AllerTop 2.0, and ToxinPred, only IFNγ inducing antigenic non-allergenic non-toxic CD4 epitopes were selected. Our analysis yielded 7 common S CD4+ T cell epitopes across Omicron (BA.1/B.1.1.529) and six other SARS-CoV-2 variants (Alpha, Beta, Delta, Gamma, US variants (S protein mutations), and Cluster 5 mink variants) reported by our other study (**Table 3, Supplementary Table S4**) [**43**]. In addition, we identified 15 Omicron BA.1 specific S CD4+ T cell epitopes **(Supplementary Table S5**), whereas 11 of them were predicted to be stable in vivo (**Table 4**). We also predicted CD4+ T cell epitopes from the BA.2 variant to check for any common epitopes among Omicron BA.1 specific CD4 peptides, and across other SARS-CoV-2 strains. Our analysis resulted in 8 common CD4+ T cell epitopes across Omicron BA.1 and BA.2 sub-lineages (**Tables 4**,**5**). Moreover, BA.2 shares identical CD4 peptides with other SARS-CoV-2 variants listed in **Table 3**.

**Table 3.**
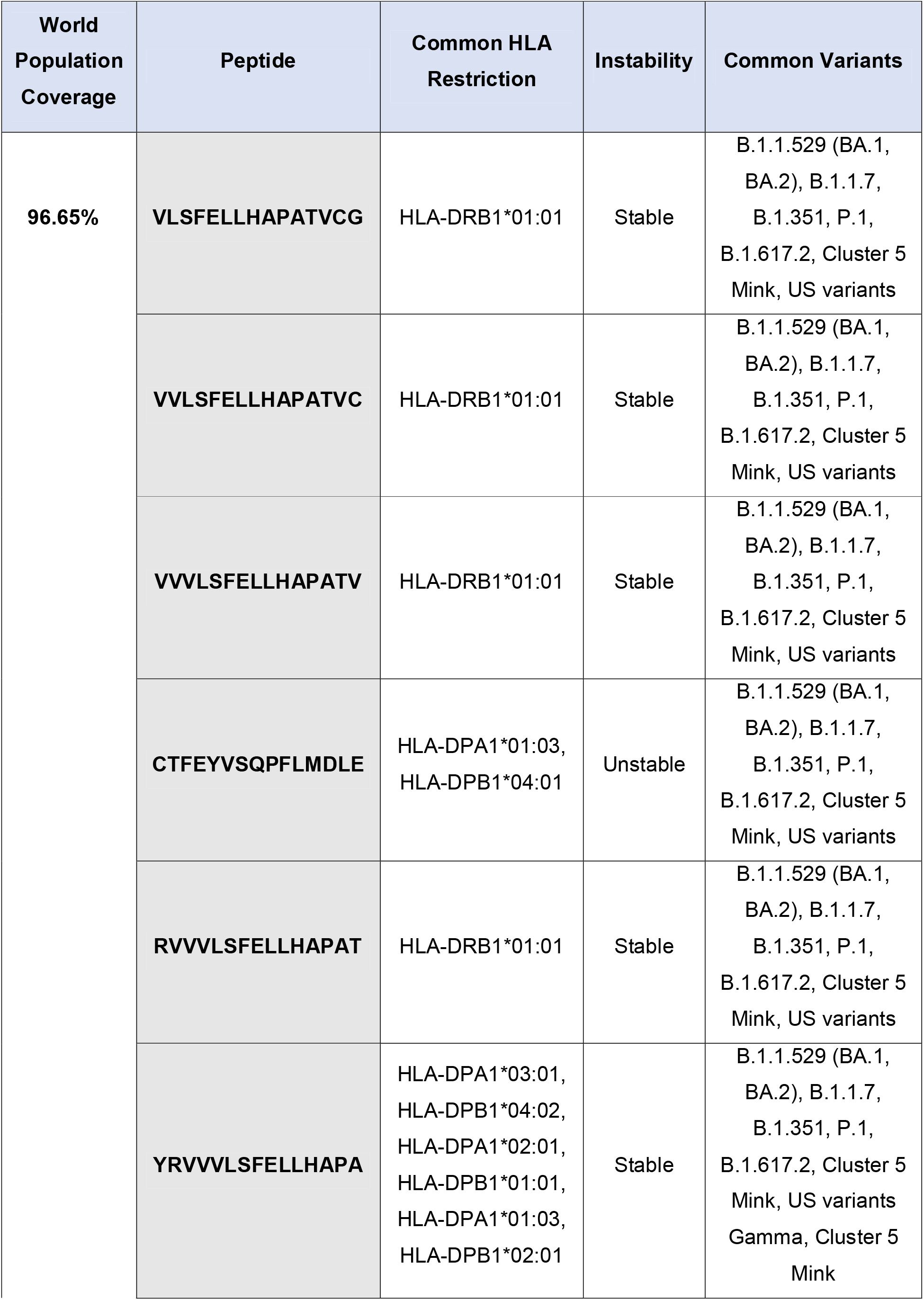
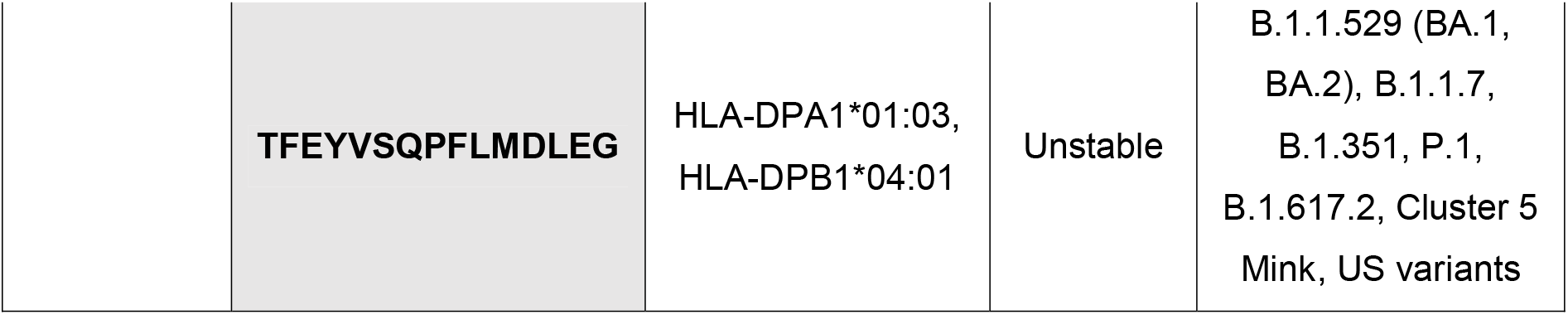
Common S protein CD4+ T cell epitopes among Omicron sub-lineages BA.1 and BA.2 as well as other SARS-CoV-2 variants including Alpha (B.1.1.7), Beta (B.1.351), Gamma (P.1), Delta (B.1.617.2), US variants (S protein mutations), and Cluster 5 mink variants. These epitopes are IFNγ inducing, antigenic, non-allergenic, and non-toxic. The world population coverage of the selected epitopes is also presented.

**Table 4.**
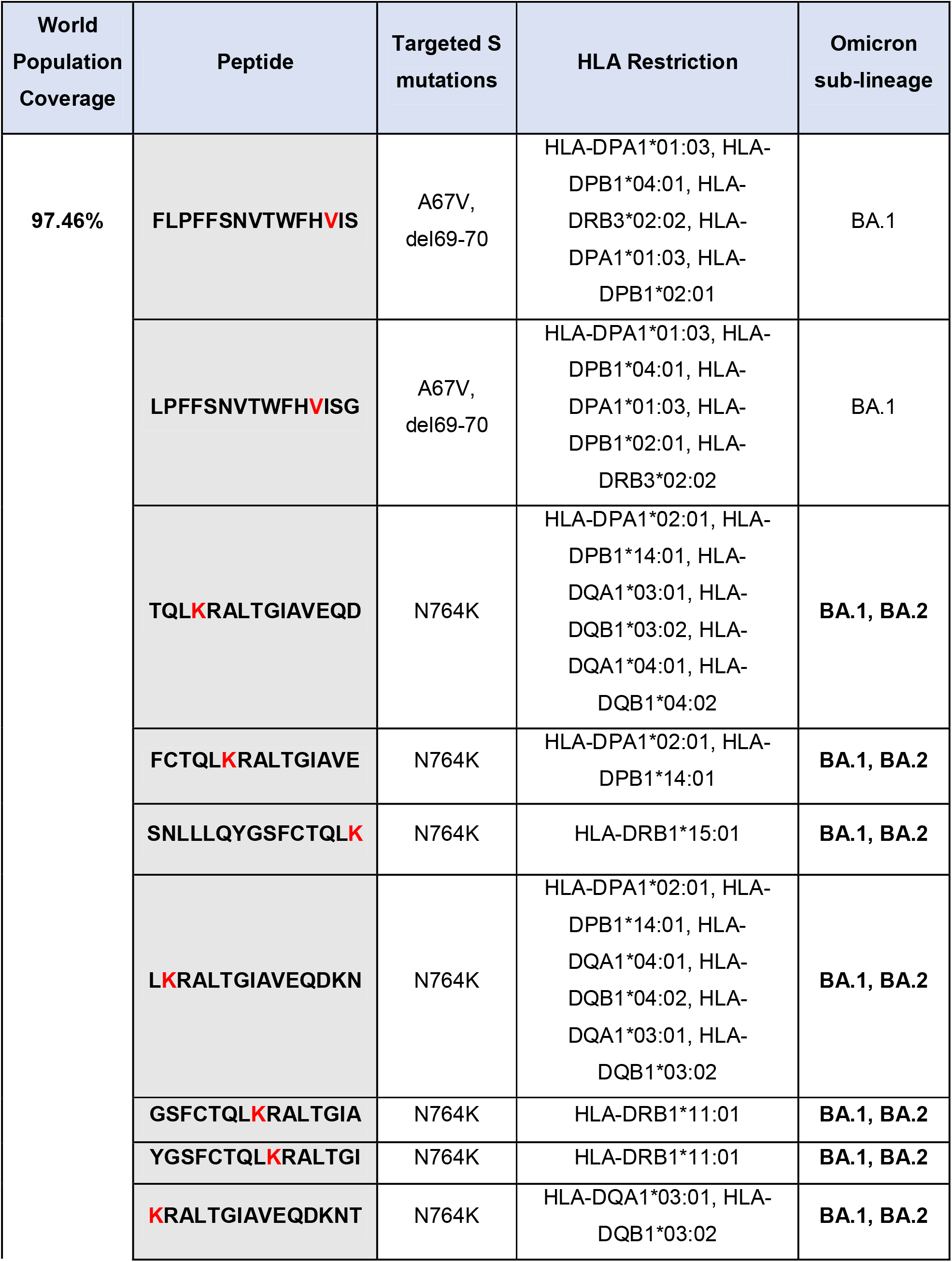
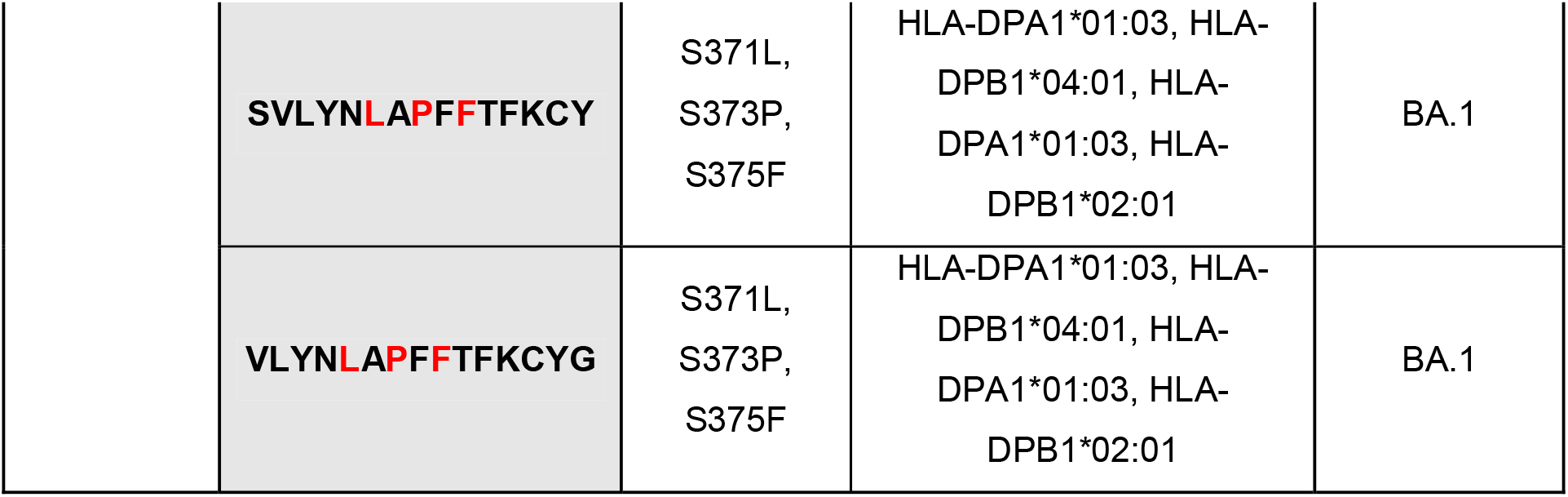
Top Omicron (BA.1/B.1.1.529) specific S protein CD4 peptides and their population coverage. All epitopes are IFNγ inducing, antigenic, non-allergenic, non-toxic, and stable. Variant-specific mutations are written in red color. The world population coverage of the selected epitopes is also presented. We identified seven common CD4+ T cell epitopes across Omicron BA.1 and BA.2 sub-lineages.

**Table 5.** Omicron (BA.1/B.1.1.529) specific IFNγ inducing, antigenic, non-allergenic, and non-toxic CD4 peptides which consist of immunogenic non-allergenic antigenic non-toxic CD8 peptides. The world population coverage of the selected epitopes is also presented. Variant-specific mutations are written in red color. We identified one overlapping CD4 and CD8 peptide (KSHRRARSVASQSII) across Omicron BA.1 and BA.2 sub-lineages.

| World Population Coverage | Omicron Specific Overlapping CD4 and CD8 Peptide | Targeted S mutations | HLA Restriction | Instability | Omicron sub-lineage |
| --- | --- | --- | --- | --- | --- |
| 92.66% | SVLYN**LAP**FFTFKCY | S371L, S373P, S375F | HLA-DPA1*01:03, HLA-DPB1*04:01, HLA-DPA1*01:03, HLA-DPB1*02:01 | Stable | BA.1 |
|  | VLYN**LAP**FFTFKCYG | S371L, S373P, S375F | HLA-DPA1*01:03, HLA-DPB1*04:01, HLA-DPA1*01:03, HLA-DPB1*02:01 | Stable | BA.1 |
|  | **KSH**RRARSVASQSII | N679K, P681H | HLA-DPA1*02:01, HLA-DPB1*14:01 | Unstable | **BA.1, BA.2** |

**Table 6.**
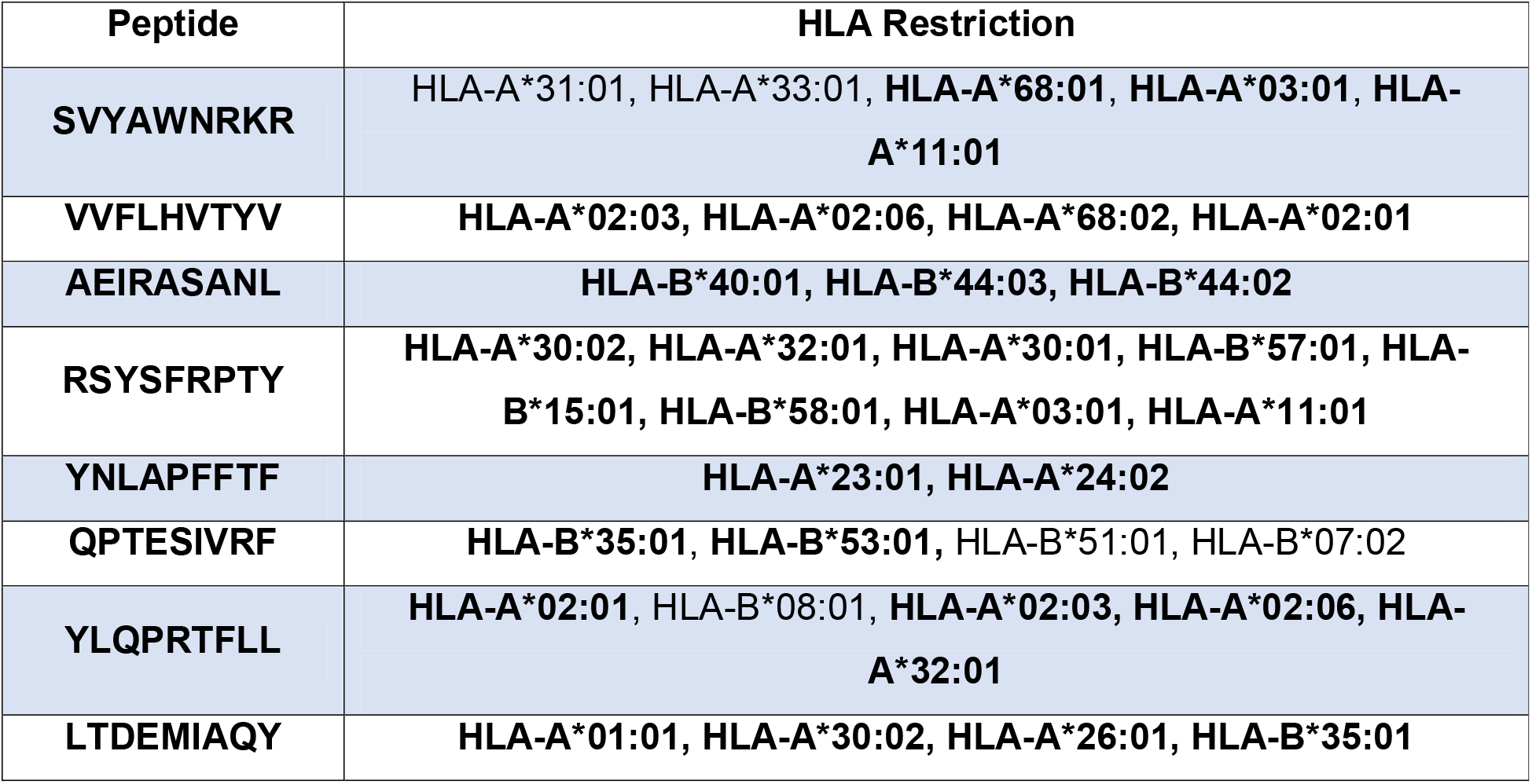
Optimized list of epitopes returned by PCOptim. Input data included all raw epitopes with a rank less than 0.30 and immunogenicity greater than 0. HLA alleles that are bolded are also present in the HLA restrictions of epitopes that passed all clinical checkpoint parameters. RSYSFRPTY and YNLAPFFTF are present in Omicronvariant-specific S CD8 epitopes and VVFLHVTYV is present in the Top common epitopes across Omicron and other SARS-CoV-2 variants.

### Identification of overlapping T cell epitopes

CD8+ T cells are important for the clearance of infected cells and CD4+ T cells play a crucial role in the promotion of specific B cell antibody production. It is thought that together these T cells can elicit a strong immune response against an evading pathogen. Hence, we overlapped the CD8+ T cell epitopes on the CD4+ T cell epitopes for the identification of overlapping Omicron BA.1 specific epitopes. The analysis from the previous section revealed 3 S CD4+ T cell epitopes (KSHRRARSVASQSII, SVLYNLAPFFTFKCY, and VLYNLAPFFTFKCYG) overlapping with our top CD8+ T cell epitopes (**Table 5**). All identified CD4+ T cell epitopes are antigenic, IFNγ inducing, non-allergenic, and non-toxic. Moreover, we found that CD4+ T cell (KSHRRARSVASQSII) that overlaps with a CD8+ T cell epitope (KSHRRARSV) is common to both Omicron BA.1 and BA.2 sub-lineages.

### Murine MHC restriction prediction

To assess the efficacy of the proposed vaccine candidates, we predicted their binding affinity to murine H2 alleles. Since the SYFPEITHI server is mostly validated for MHC Class I epitope prediction and has low reliability (50%) for MHC Class II peptides, we only identified CD8 peptides that bind to murine H2 alleles. The comparison of reference epitopes to the identified CD8 peptides yielded potential strong, intermediate, and weak binders. Our analysis revealed 7 common S CD8+ T cell epitopes among SARS-CoV-2 variants (**Table 1**) and 4 Omicron BA.1 specific S CD8+ T cell epitopes (**Table 2**) presented by murine MHC restriction. Moreover, we identified one BA.2 lineage-specific CD8+ T cell epitope (TPINLGRDL) that is presented by murine MHC restriction (**Supplementary Table S3**).

### Population coverage analysis

To determine the world population coverage of the selected T cell epitopes, corresponding HLA associations were considered. Identified epitopes that bind several MHC (HLA) alleles are generally regarded as the best probable epitopes as they have a higher potential to show good coverage by approaching 100%. Subsequently, we found that the 12 common CD8 peptides among Omicron and other SARS-CoV-2 variants cover 84.0% of the world population (**Table 1, Supplementary Fig. 1A**) whereas the 8 Omicron BA.1 specific CD8 peptides cover 76.16% (**Table 2, Supplementary Fig. 1B**). World population coverage was also computed for the MHC Class II peptides. We found that the 7 common CD4+ T cell epitopes across Omicron and other circulating variants including Delta can elicit an immune response that covers 96.65% of the world population (**Table 3, Supplementary Fig. 1C**). The 11 top quality Omicron BA.1 variant-specific CD4 peptides were computed to cover 97.46% of the world population (**Table 4, Supplementary Fig. 1D**) whereas the three overlapping CD4 peptides were found to cover 92.66% of the world population (**Table 5, Supplementary Fig. 1E**).

### Validation of top epitopes

The top 300 epitopes found from IEDB NetMHCpan EL 4.1 have a rank range of 0-0.30. We filtered through all the raw epitope data to obtain a dataset of epitopes with a rank less than 0.30 and immunogenicity greater than 0. We entered this dataset into PCOptim to obtain a list of optimized epitopes which we can use as a basis of comparison to our 8 immunogenic, antigenic, non-allergenic, non-toxic, and stable epitopes. The optimized list of epitopes included 8 unique epitopes and returned 98.55% population coverage (**Figure 6**). Of the 8 optimized epitopes, 2 are present in our “Omicron BA.1 variant-specific S CD8 epitope” dataset (**Supplementary Table S2)** (RSYSFRPTY and YNLAPFFTF) and 1 is present in our “Top common S CD8 epitopes across Omicron and other SARS-CoV-2 variants” dataset (**Supplementary Table S1**) (VVFLHVTYV). The HLA alleles that were present in the optimized dataset but not the clinically relevant dataset are HLA-B*07:02, HLA-B*08:01, HLA-A*31:01, HLA-A*33:01, and HLA-B*51:01. These alleles account for the difference in population coverage between the optimal epitope dataset and datasets obtained after passing epitopes through several clinical checkpoints.

### Three-dimensional (3D) structural analysis

Protein 3D structures offer useful insights into their molecular activity and provide a wide variety of applications in bioscience. We selected two immunogenic antigenic non-allergenic and non-toxic CD8+ T cell epitopes (KSHRRARSV and YNLAPFFTF) for 3D visualization and molecular docking analysis. KSHRRARSV and YNLAPFFTF are both strong MHC I binders and presented by murine MHC restriction. Epitope KSHRRARSV was docked with HLA-A*30:01 (**Figure 7A**) and epitope YNLAPFFTF with HLA-A*24:02 (**Figure 7B**). The structures were also energy minimized to reduce the overall potential energy between the epitope and MHC molecule. Furthermore, to get a better insight into the TCR interaction with the pMHC complex, we modeled the YNLAPFFTF-HLA-A*24:02 complex with the C1-28 TCR (RCSB PDB: 3VXM) structure specific to HLA-A24 (**Figure 7C**).

## Discussion

Although several COVID-19 vaccines are currently available, the excessive mutations observed in the spike protein of Omicron can escape immune response, raising concerns over the efficacy of these vaccines [**44**,**46**]. Despite the overall success of mRNA vaccines which have provided a breakthrough technology in protecting the community against the severity of infection, it remains urgent to expand the landscape of vaccines to combat the novel Omicron variant as well as any future SARS-CoV-2 strains [**47**,**48**]. Here, we present an epitope-based vaccine design that has many advantages over the current mRNA vaccines. Epitope-based vaccines are highly specific, having the ability to target multiple SARS-CoV-2 antigens simultaneously. They also confer a long-lasting immune response, superior to the relatively short-lived mRNA vaccines which have shown to decline after 6 months of vaccination [**49**]. Another important advantage of epitope-based vaccines is their absence of adverse side effects which avoid allergenic consequences.

To design an efficient and safe vaccine that would provide broad protection across many ethnicities, we carefully selected the prediction tools based on their accuracy. The IEDB prediction servers are free and widely accepted in literature [**50**]. However, to select the most reliable T cell prediction tool, we used our previous study that analyzed the specificity and sensitivity of several MHCI and MHCII peptide prediction tools [**43**]. We used NetMHCpan EL 4.1 for CD8 epitope predictions and IEDB Recommended 2.22 for CD4 epitope predictions. Due to the lack of reliable immunogenicity prediction tools for CD4 peptides on IEDB.org, the IFNEpitope server was used to predict IFNγ inducing epitopes. IFNEpitope is predominantly designed to predict MHCII peptides with the capacity to induce IFNγ release and has an accuracy of 82.10%. The VaxiJen and AllerTOP servers have been regarded as highly reliable antigenicity and allergenicity prediction servers with an accuracy of 87% and 89%, respectively. For toxicity evaluation, ToxinPred was selected due to its accuracy of 94.50%.

We revealed CD8+ and CD4+ T cell epitopes from SARS-CoV-2 S protein targeting the Omicron (BA.1/B.1.1.529) variant while considering many of its significant mutations such as A67V, del69-70, T95I, ins214EPE, S371L, S373P, S375F, Q493K, G496S, Q498R, N501Y, N679K, and P681H. We identified 8 immunogenic antigenic non-allergenic non-toxic CD8+ T cell epitopes (**Figure 3**) and 11 IFNγ inducing antigenic non-allergenic non-toxic stable CD4+ T cell epitopes that can provide a robust immune response and cover 76.16% and 97.46% of the world population, respectively (**Tables 1 and 2**). Among the identified epitopes, 5 CD8 peptides (RSYSFRPTY, NLAPFFTFK, YNLAPFFTF, APFFTFKCY, and VLYNLAPFF) are located on the RBD region, suggesting their potential to target the impaired interaction between S protein and the neutralizing antibodies. To give a better insight into how our epitopes target Omicron S protein RBD mutations, we obtained the crystal structure of Fab fragment in complex with SARS-CoV-2 S protein (RCSB PDB 6XCM) (**Figure 4**). Moreover, our predicted CD8 epitope RSYSFRPTY targets the two important mutations, Q498R and N501Y, correlated with an enhanced binding affinity to the ACE2 and decreased binding affinity to neutralizing antibodies [**51**]. We predicted RSYSFRPTY to be immunogenic, highly antigenic, non-allergenic, and non-toxic with a world population coverage of 49.04% alone, suggesting its importance in epitope-based vaccine design. To further enhance the immune response, we identified two 15-mer peptides (SVLYNLAPFFTFKCY, VLYNLAPFFTFKCYG) that overlap with the identified CD8+ T cell epitopes on the RBD region. Together these epitopes can provide a strong and long-lasting immune response against the Omicron variant.

**Figure 3.**
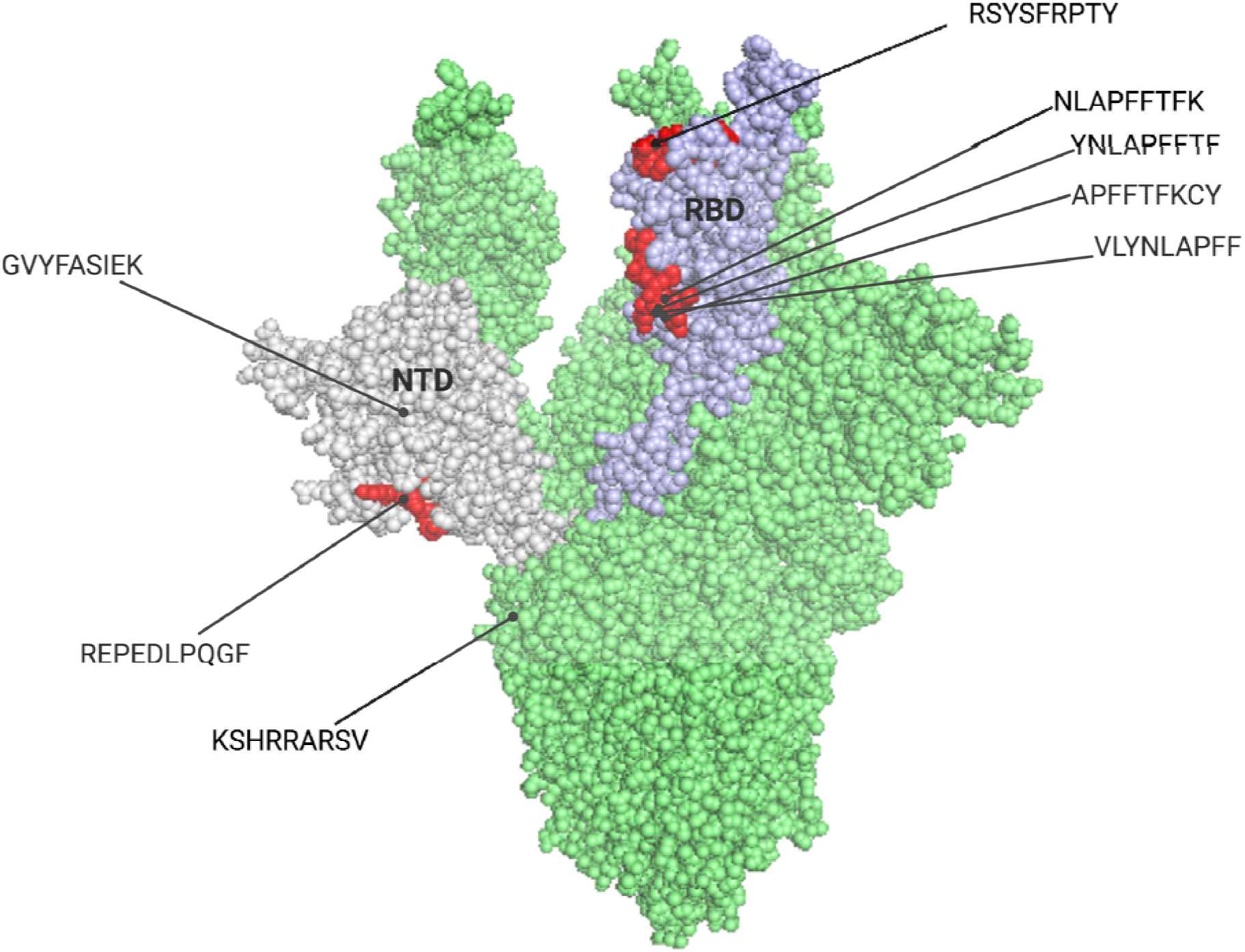
3D structure of SARS-CoV-2 S protein (PDB: 7VNE) with highlighted CD8+ T cell epitopes targeting newly evolved acquired mutations in Omicron (BA.1/B.1.1.529) variant. These epitopes are located on the RBD, NTD, and furin cleavage sites and can be used as part of the multi-epitope vaccine. All epitopes were predicted to be immunogenic, antigenic, non-allergenic, and non-toxic.

**Figure 4.**
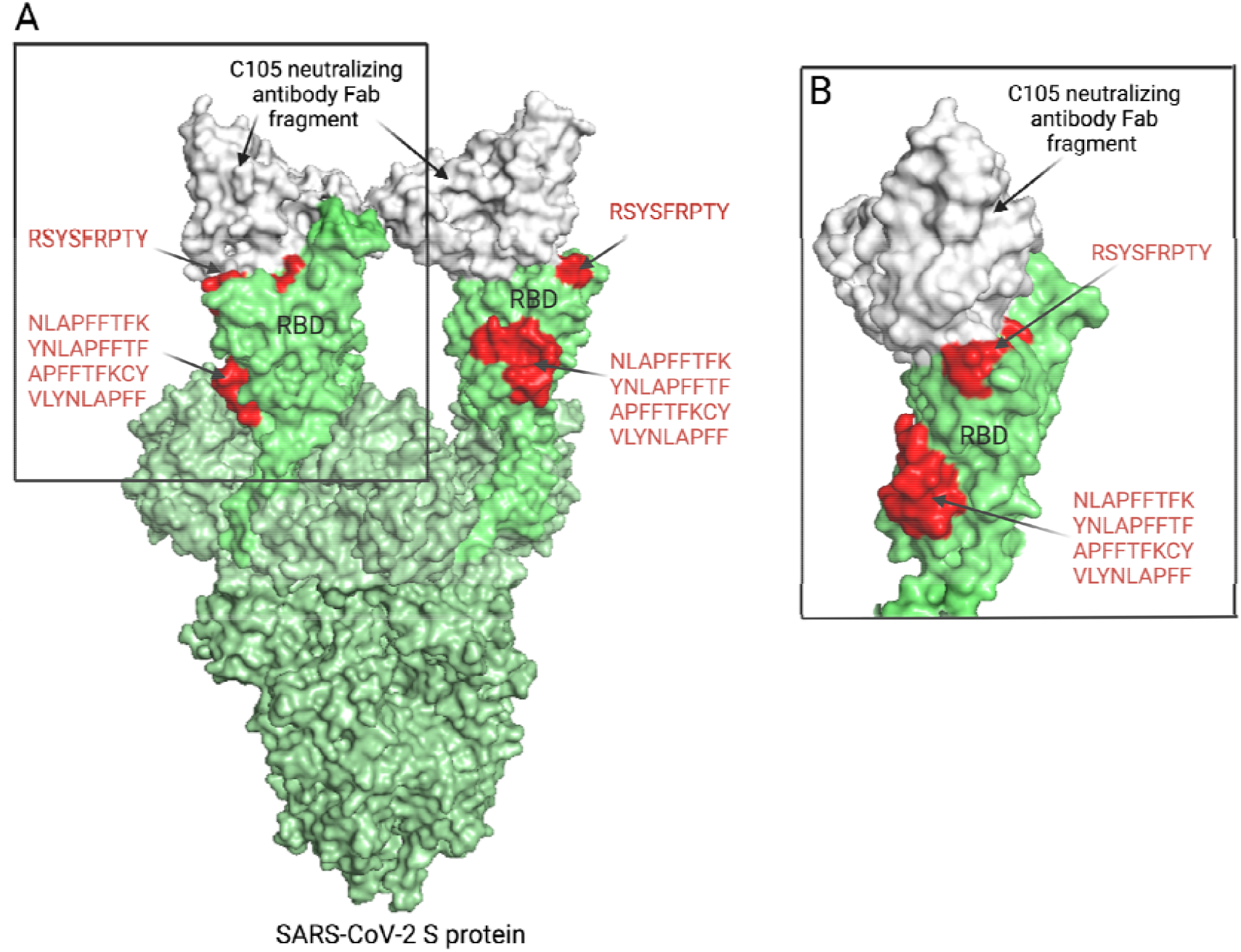
3D structure of SARS-CoV-2 S protein in complex with the C105 neutralizi antibody Fab fragment (PDB: 6XCM) with five highlighted CD8+ T cell epitopes specific for Omicron (BA.1/B.1.1.529) variant. (**A**) SARS-CoV-2 S protein in complex with the C105 neutralizing antibody Fab fragment. Our predicted immunogenic CD8+ T cell epitopes (RSYSFRPTY, NLAPFFTFK, YNLAPFFTF, APFFTFKCY, and VLYNLAPFF) are colored in red. (**B**) Zoomed in version of the complex that focuses on S protein RBD interaction with the neutralizing antibody, highlighting the predicted epitopes.

We used our software tool PCOptim to validate the accuracy of the CD8 top epitope selection. Additionally, the PCOptim tool allowed us to determine the caveats in population coverage and thus identify the populations that may receive less coverage. The PCOptim tool provided us with an optimal set of epitopes that reaches the maximum possible population coverage given our raw dataset. Comparing this optimal dataset to our set of top epitopes revealed some overlap, confirming the success of our selection procedure. A total of three epitopes were found in both the optimal dataset and our top epitope datasets. Out of the 27 most common HLA alleles in the human population, we identified 5 alleles that are unaccounted for in the immunogenic, antigenic, non-allergenic, non-toxic, and stable epitope dataset (HLA-B*07:02, HLA-B*08:01, HLA-A*31:01, HLA-A*33:01, and HLA-B*51:01). The difference in population coverage between that optimal dataset and our set of 8 CD8 Omicron-related epitopes can be attributed to the five aforementioned HLA alleles. The only way to increase population coverage further is to include new epitopes that are predicted to be strong binders to the desired HLA alleles and pass the clinical checkpoint filters. While PCOptim does not account for the clinical checkpoint parameters we address in this study, it is a useful tool in determining several epitopes that are useful in obtaining high population coverage. The population coverage that we obtained from our top epitope dataset is substantially high, so adding new epitopes will make a very small impact on total population coverage.

Previous studies have shown that SARS-CoV-2 may use mutations on the NTD region of S protein to escape potent polyclonal neutralizing responses [**52**]. Subsequently, we identified an immunogenic antigenic non-allergenic and non-toxic CD8+ T cell epitope REPEDLPQGF which consists of a novel insertion mutation ins214EPE, not observed in any other SARS-CoV-2 strains [**53**]. Although the mutation is believed to arise from co-infection of SARS-CoV-2 with HCoV-229E as the two share similar nucleotide sequences, we were unable to detect any common epitopes between the two human-coronavirus strains after running a multiple sequence alignment analysis. To create a robust immune response that targets the NTD region of SARS-CoV-2 Omicron, we identified an additional CD8+ T cell epitope (GVYFASIEK) and two high-quality CD4+ T cell epitopes (FLPFFSNVTWFHVIS and LPFFSNVTWFHVISG) which consist of mutation A67V and a pair of deletions at residues 69 and 70.

We predicted another important immunogenic antigenic non-allergenic non-toxic CD8 peptide (KSHRRARSV) which is located at the furin cleavage site of SARS-CoV-2. The cleavage of the S protein into S1 and S2 is an essential step in viral entry into a host cell [**54**] and it is believed that the mediation of membrane fusion could be linked to the high virulence of SARS-CoV-2, mainly due to a cluster of mutations at the H655Y, N679K and P681H amino acid sites [**18**]. KSHRRARSV targets both N679K and P681H and is presented by an antigenic IFNγ inducing non-allergenic non-toxic 15-mer amino acid sequence (KSHRRARSVASQSII) which together can provide strong protection against B.1.1.529 or any future variants. These epitopes also target the novel circulating sister-lineage BA.2 of Omicron which confers identical mutations observed in B.1.1.529.

A large number of mutations on the RBD of S protein is speculated to help the virus escape neutralizing antibodies from natural and vaccine-induced immunity. Despite the loss of binding affinity to human ACE2 due to mutations such as K417N, Omicron has restored its strong binding to the ACE2 with mutations at residues 493, 496, 498, and 501. Previously we showed how epitopes can target the S protein interaction with neutralizing antibodies (**Figure 4**). While completing the manuscript, Omicron-specific S protein structure in complex with human ACE2 (PDB 7T9K) became available which we used to visualize our predicted epitopes and their interaction with human ACE2 and Omicron S protein complex. We show that CD8+ T cell epitopes (RSYSFRPTY, NLAPFFTFK, YNLAPFFTF, APFFTFKCY, and VLYNLAPFF) on the RBD target the ACE2 binding site (**Figure 5**). Together with the predicted CD4+ T cell epitopes, a robust immune response can be created which facilitates the production of antibodies, therefore blocking cell entry. Moreover, NTD and furin cleavage sites are critical to virus attachment to membrane protein and other sites, and we have identified epitopes that can target these sites (**Figure 5**). With the emergence of new SARS-CoV-2 variants and their high likelihood to adapt to mutations on the RBD, NTD, and furin cleavage sites, we speculate that our identified epitopes can effectively target any future variants by blocking their binding to human ACE2. We have already observed that the BA.2 lineage confers many common mutations in the BA.1/B.1.1.529 variant. We show that CD8 peptide (KSHRRARSV) and CD4 epitope (KSHRRARSVASQSII) are recognized by both Omicron sub-lineages (BA.1 and BA.2) (**Tables 2**,**5**). Moreover, these epitopes are located on the S1/S2 cleavage site, associated with increased ACE2 binding affinity and impaired antibody recognition. Therefore, we propose that these epitopes can be used to target the critical site of SARS-CoV-2 S protein in both BA.1 and BA.2 lineages.

**Figure 5.**
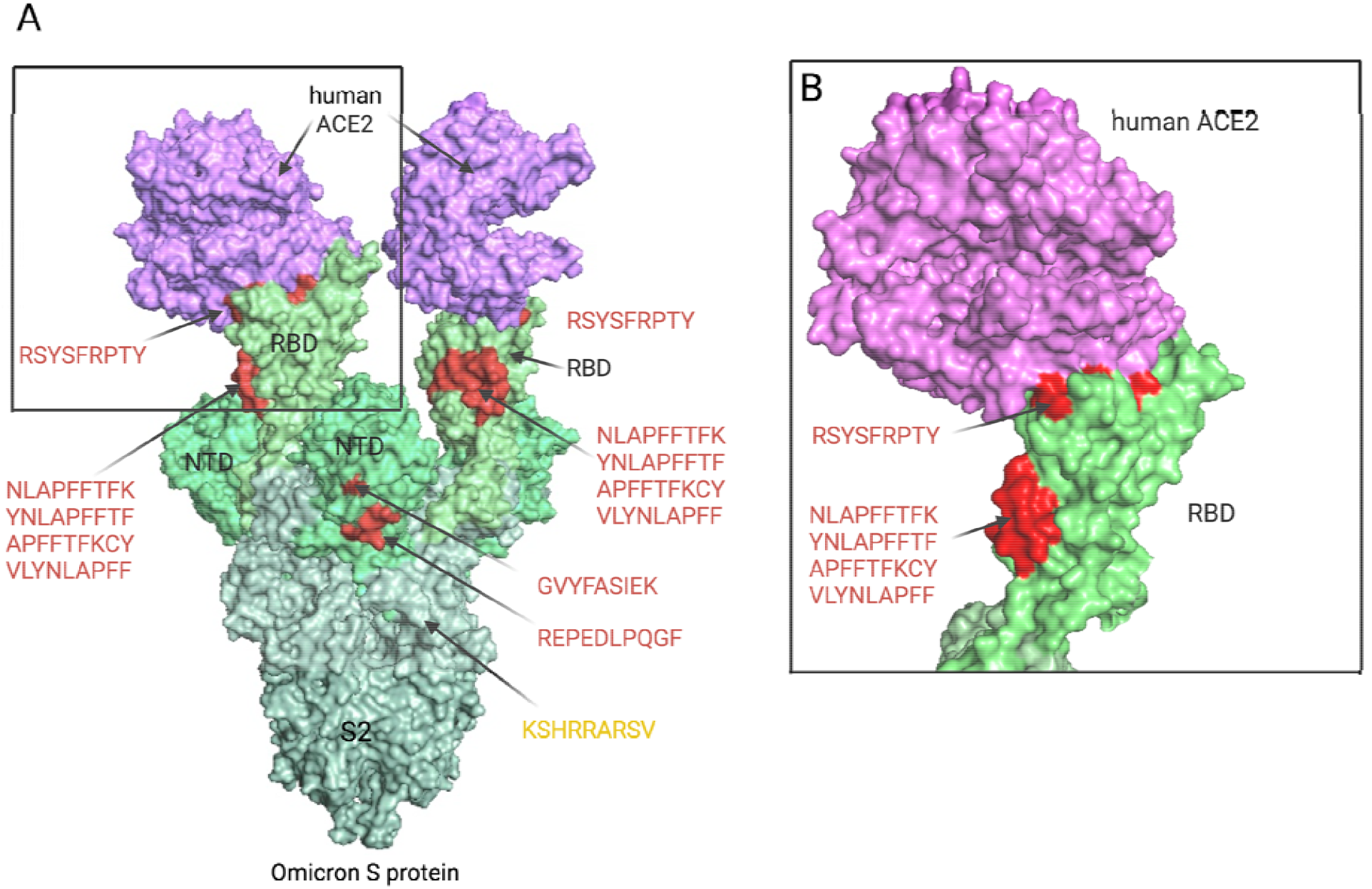
3D structure of SARS-CoV-2 Omicron S protein-ACE2 complex (PDB: 7T9K) with eight highlighted CD8+ T cell epitopes. Original Omicron (BA.1/B.1.1.529) specific epitopes are colored in red. Common epitopes among BA.1 and BA.2 variants are colored in yellow. (**A**) Omicron S protein in complex with human ACE2. Five of our predicted immunogenic CD8+ T cell epitopes (RSYSFRPTY, NLAPFFTFK, YNLAPFFTF, APFFTFKCY, and VLYNLAPFF) are within the receptor-binding domain (RBD), targeting the binding site of ACE2. Two immunogenic CD8+ T cell epitopes (GVYFASIEK and REPEDLPQGF) target novel Omicron (BA.1) specific mutations on the NTD. One immunogenic CD8+ T cell epitope (KSHRRARSV) was identified at the furin cleavage site, common for both BA.1 and BA.2 lineages. (**B**) Zoomed in version of the complex that focuses on Omicron RBD interaction with human ACE2, highlighting the predicted CD8 epitopes on the RBD site.

**Figure 6.**
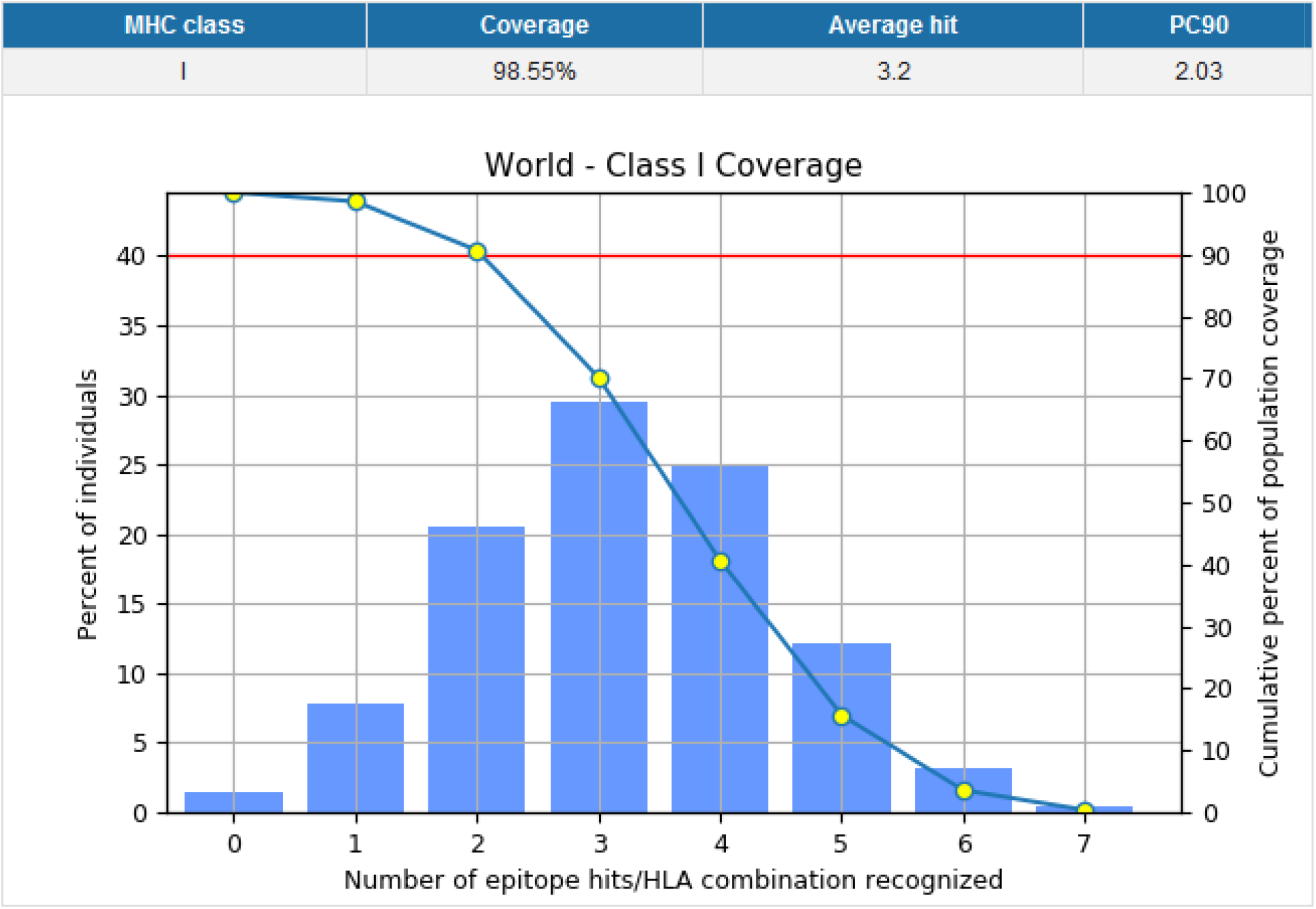
Population coverage for optimal epitope dataset retrieved from our PCOptim software tool. The epitope data accounts for 27 unique HLA alleles which contribute to the high population coverage. Epitope datasets that pass through the clinical checkpoint filters will have fewer unique HLA alleles and therefore have lower population coverage.

**Figure 7.**
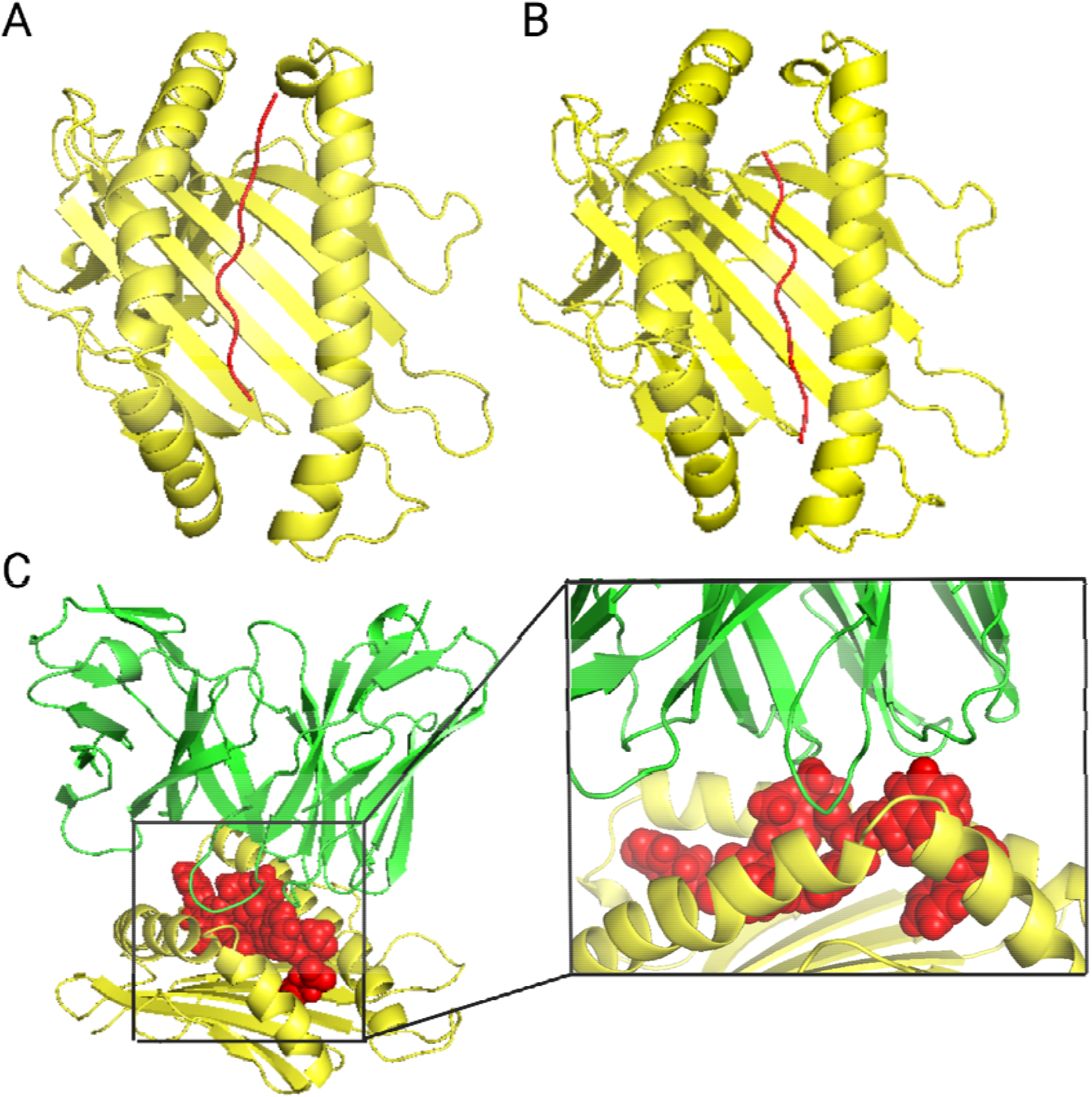
3D modeling of two SARS-CoV-2 S peptides in complex with corresponding MHC Class I molecule. **(A)** Cartoon form of CD8+ T cell epitope KSHRRARSV (red) and HLA-A*30:01 allele (yellow). **(B)** Cartoon form of CD8+ T cell epitope YNLAPFFTF (red) and HLA-A*24:02 allele (yellow). **(C)** Binding interaction of sphere form of epitope YNLAPFFTF (red) with C1-28 TCR (green) (PDB: 3VXM) and HLA-A*24:02 (yellow).

As it is highly important to design an effective booster shot against the novel SARS-CoV-2 Omicron variant, we also attempted to design a protective vaccine against other SARS-CoV-2 strains. Subsequently, we identified 12 immunogenic CD8+ and 7 IFNγ inducing CD4+ T cell epitopes across Alpha, Beta, Delta, Gamma, Omicron (BA.1, BA.2), US variants (S protein mutations), and Cluster 5 mink variants that are predicted to protect 84.0% and 96.65% of the world population, respectively (**Tables 1 and 3**). Moreover, all of the CD8 and CD4 epitopes were predicted to be antigenic, non-allergenic, and non-toxic for safe vaccine development. These epitopes will likely reactivate B and T cells against the original epitopes without creating a variant-specific immune response.

The findings through computational analysis indicate that the designed epitopes could be vastly useful in multi-epitope vaccine construct as they have a high probability of being safe and effective in protecting humans from the SARS-CoV-2 virus. To further induce the immunogenicity of the antigens and provide a more robust and long-lasting immune response, we suggest that emulsion adjuvant, like MF59, could be added to the final vaccine construct. Superior to other types of adjuvants, they elicit a cell-mediated immune response with improved antigen uptake and the ability to promote the migration of activated antigen-presenting cells. Furthermore, MF59 has been used in preclinical trials of coronavirus targeted vaccines [**55**]. Linkers are also important for vaccines while providing enhanced flexibility, extended protein folding, and separation of functional domains. Available studies propose that CD8 and CD4 epitopes could be linked together using AYY linker and GPGPG linker, respectively [**56**].

It is detrimental to assess the safety and efficacy of proposed multi-epitope vaccine candidates in animal models. Murine models have low-cost maintenance and reflect the clinical signs, viral replication, and pathology of SARS-CoV-2 in humans [**57**]. We used the SYFPEITHI prediction server to predict murine MHC binding affinity to MHC I epitopes. The prediction server utilizes published T cell epitopes and motifs and is validated to be 80% accurate. We identified four Omicron BA.1 specific and one BA.2 specific S CD8+ T cell epitopes, as well as 7 common S CD8+ T cell epitopes among SARS-CoV-2 variants presented by murine MHC restriction. Ultimately, the predicted strong, intermediate, and weak MHCI binders can be used in further preclinical trials.

### Limitations of study

The *in-silico* prediction analysis of the SARS-CoV-2 S protein reported in our study requires further laboratory validation to select the most efficient vaccine candidate.

## Conclusion

The recent emergence of the highly mutated Omicron variant of SARS-CoV-2 has disrupted confidence around whether the current vaccines and antibody therapies will provide long-term protection against the novel coronavirus. Subsequently, the possibility of escape from natural and vaccine-induced immunity has prompted an urgent need for new vaccine constructs which target the most concerning mutations on the variant. Using immunoinformatic methods and bioinformatics tools, we identified immunogenic antigenic non-allergenic non-toxic CD8+ T cell epitopes and IFNγ inducing antigenic non-allergenic non-toxic CD4+ T cell epitopes on the RBD, NTD, and furin cleavage sites of S protein which target Omicron specific mutations by blocking the binding of S protein to ACE2. Moreover, our analysis yielded common high-quality CD8+ and CD4+ T cell epitopes across Omicron (BA.1, BA.2) and other circulating SARS-CoV-2 variants including Delta, Alpha, Beta, Gamma, US variants (S protein mutations), and Cluster 5 mink variants with a world population coverage of 84.0% and 96.65%, respectively. We validated the findings using our software PCOptim which enabled us to identify an optimal set of epitopes reaching the maximum possible population coverage. Our *in-silico* epitope prediction together with murine MHC affinity prediction enables scientists to validate the efficacy of the proposed multi-peptide vaccine model through further preclinical studies. To understand the structural validation of the vaccine candidate and the binding affinity to MHC and TCR molecules, we performed a molecular docking analysis. To conclude, the multi-epitope vaccine constructs designed against S protein of SARS-CoV-2 by utilizing immunoinformatic methods may be considered as a new, safe, and efficient approach to control the Omicron variant as well as any future SARS-CoV-2 variants.

## Acknowledgments

We wish to acknowledge Lombardi Comprehensive Cancer Center, and Georgetown University Medical Center for their support.

## Notes

### Competing Interest Statement

The authors have declared no competing interest.

