## Supplementary Figure S1 for "SARS-CoV-2 Omicron (BA.1 and BA.2) specific novel CD8+ and CD4+ T cell epitopes targeting spike protein"

**A**	**B**

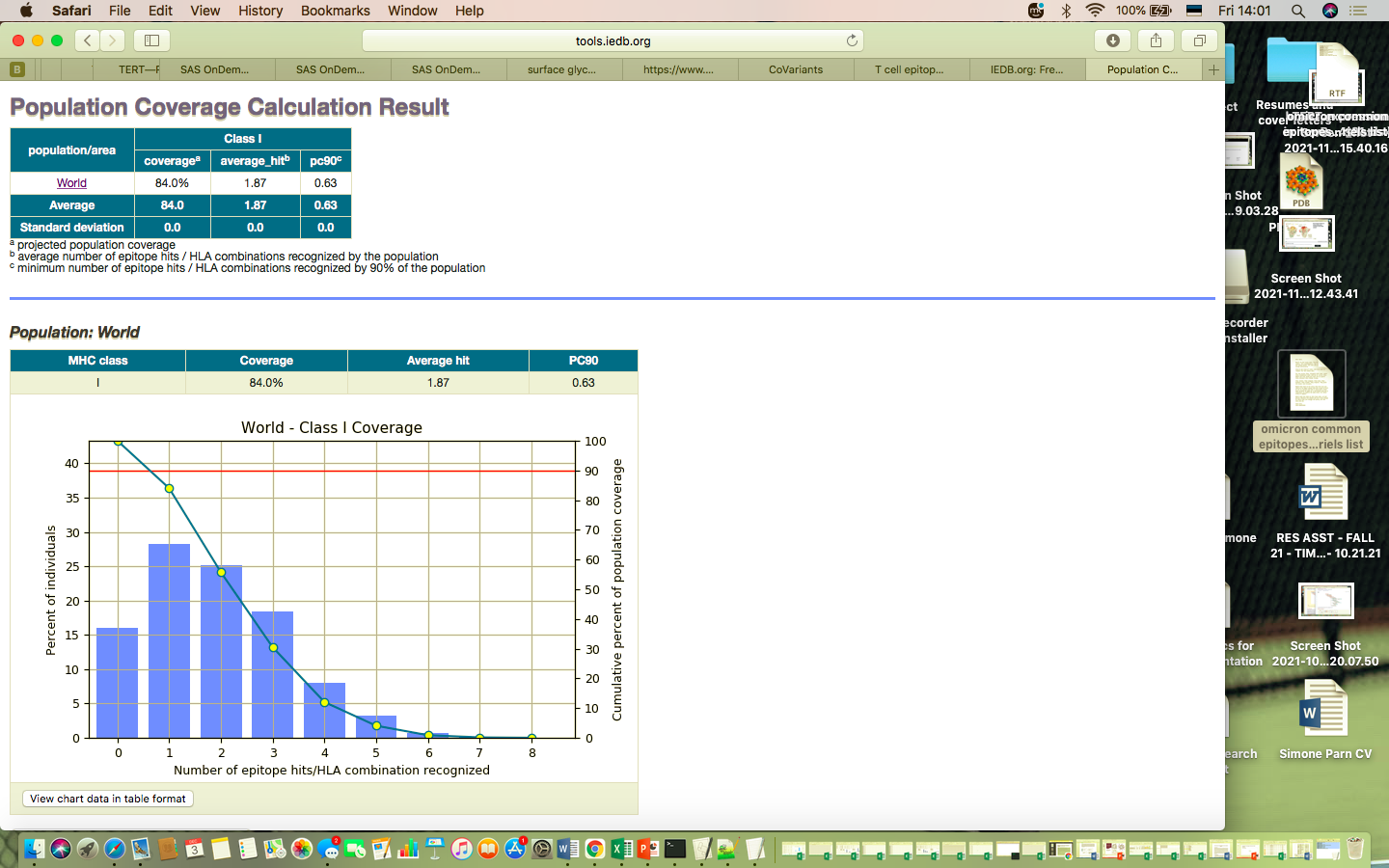

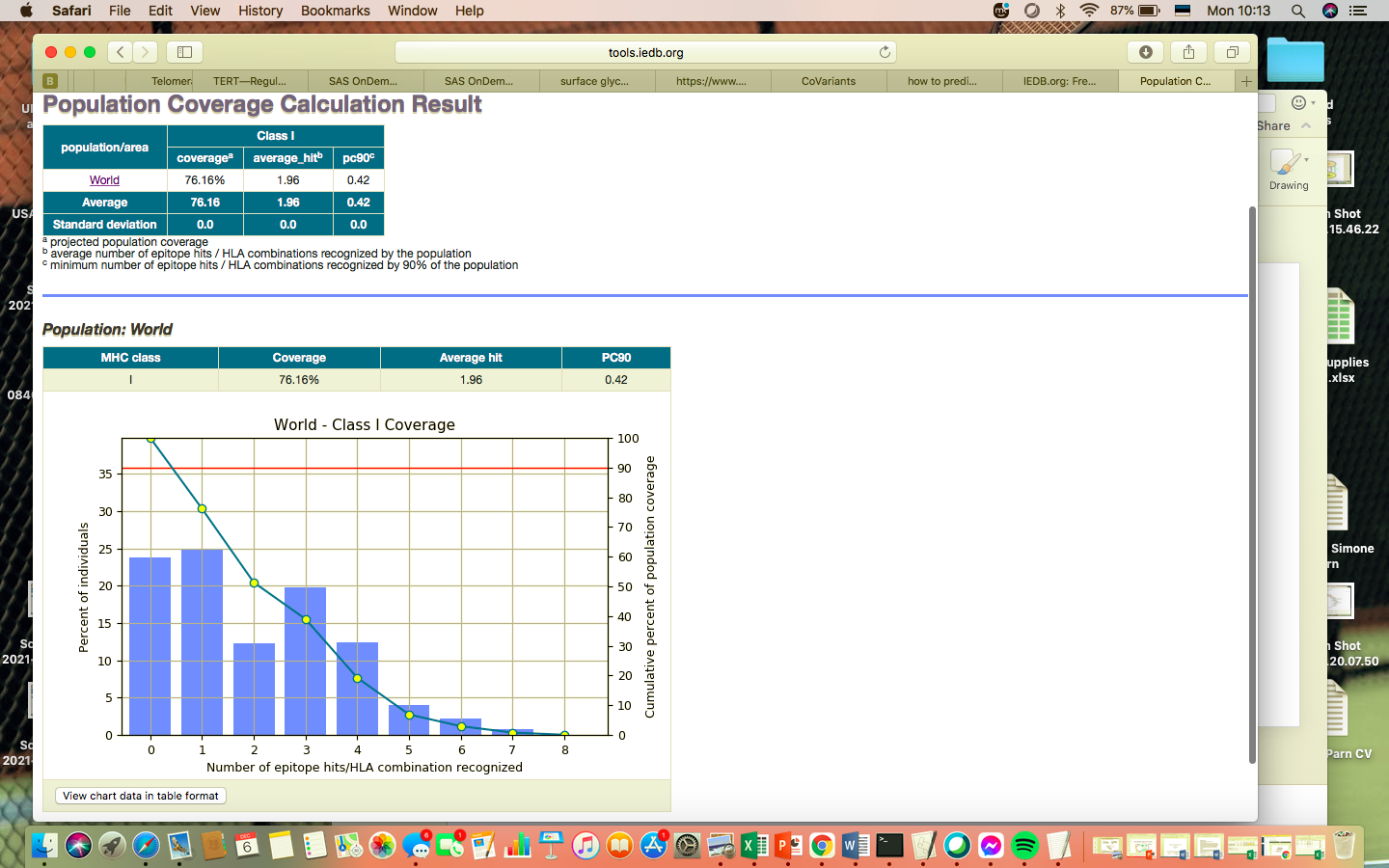

**C	D**

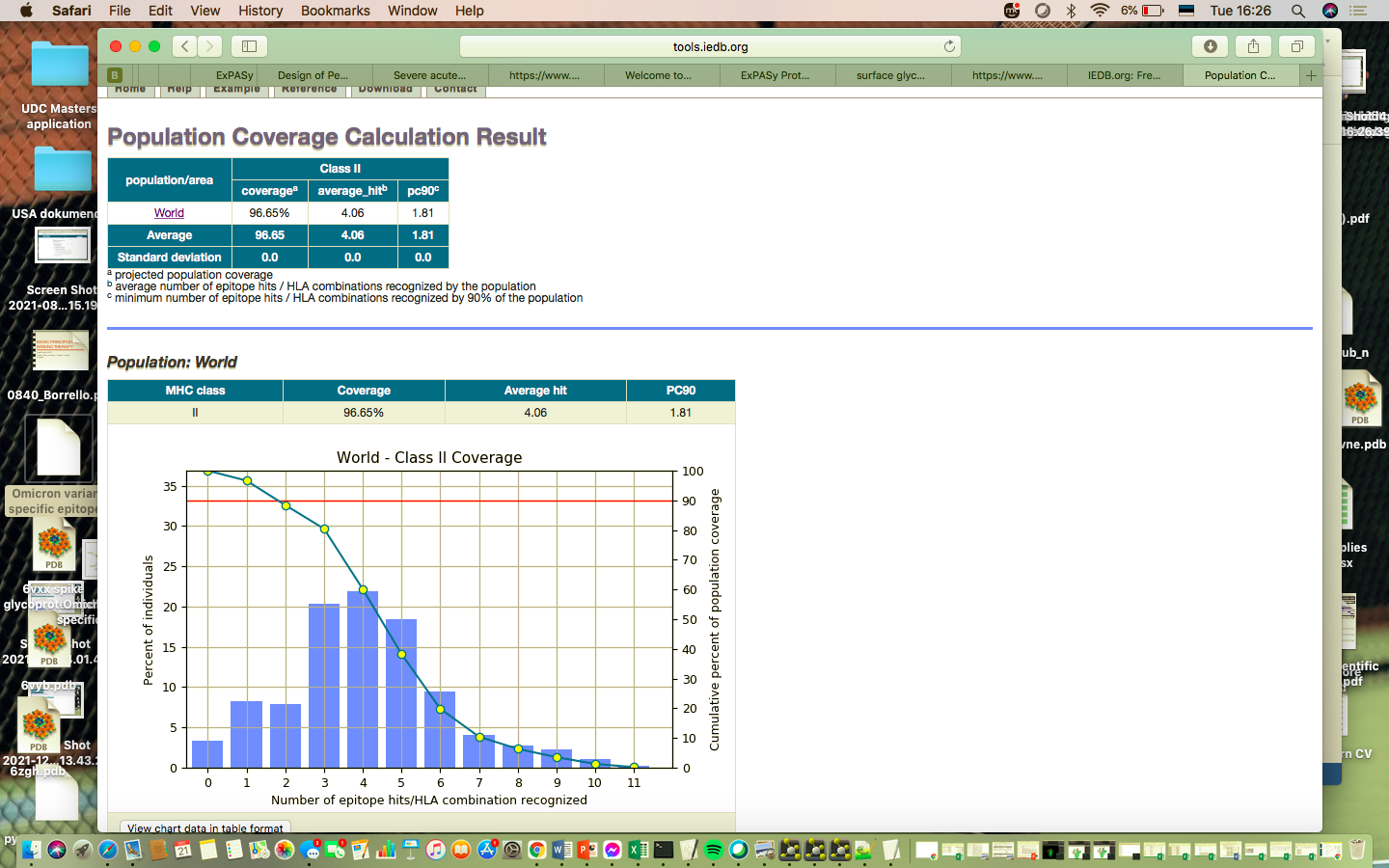

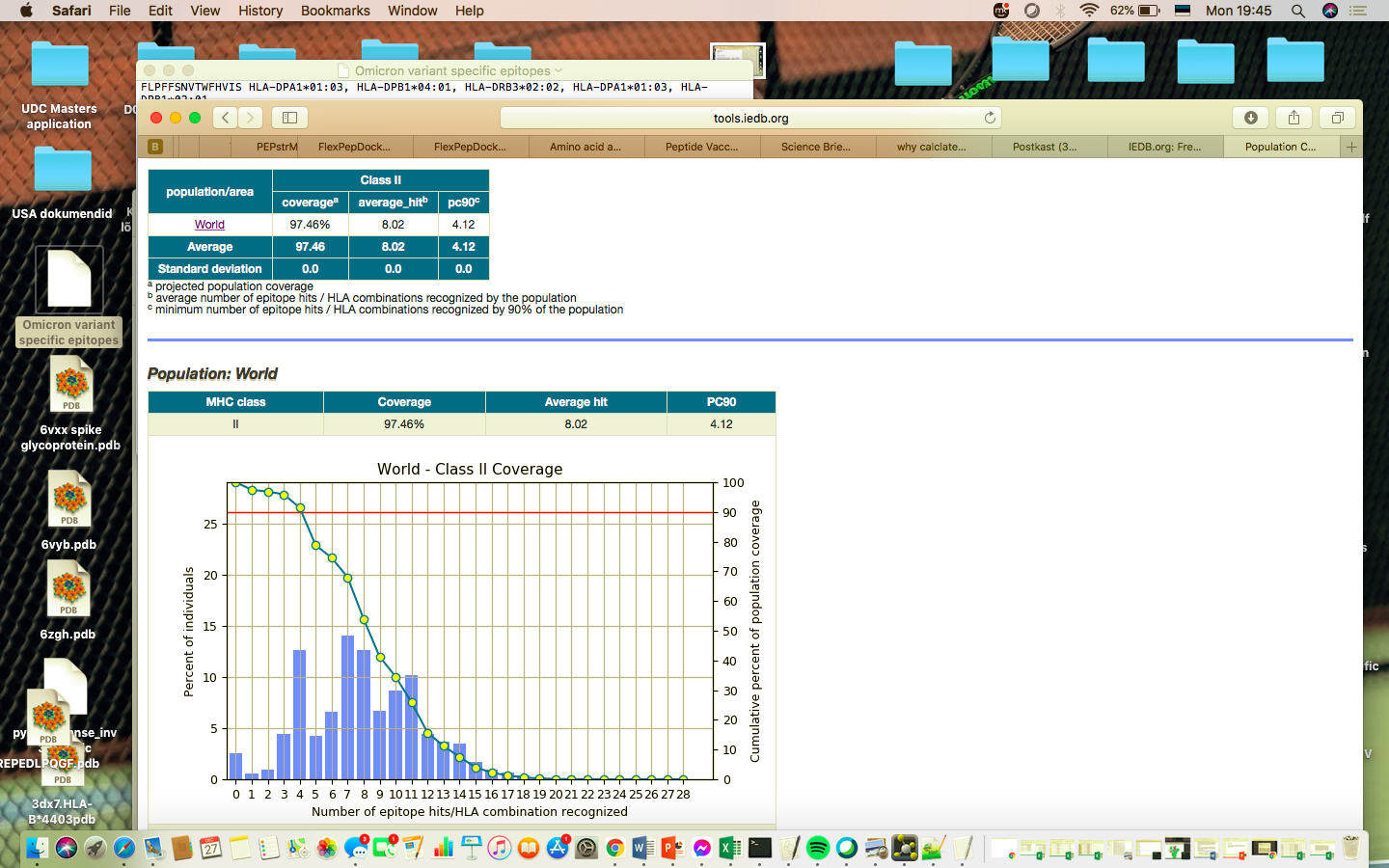

**E**

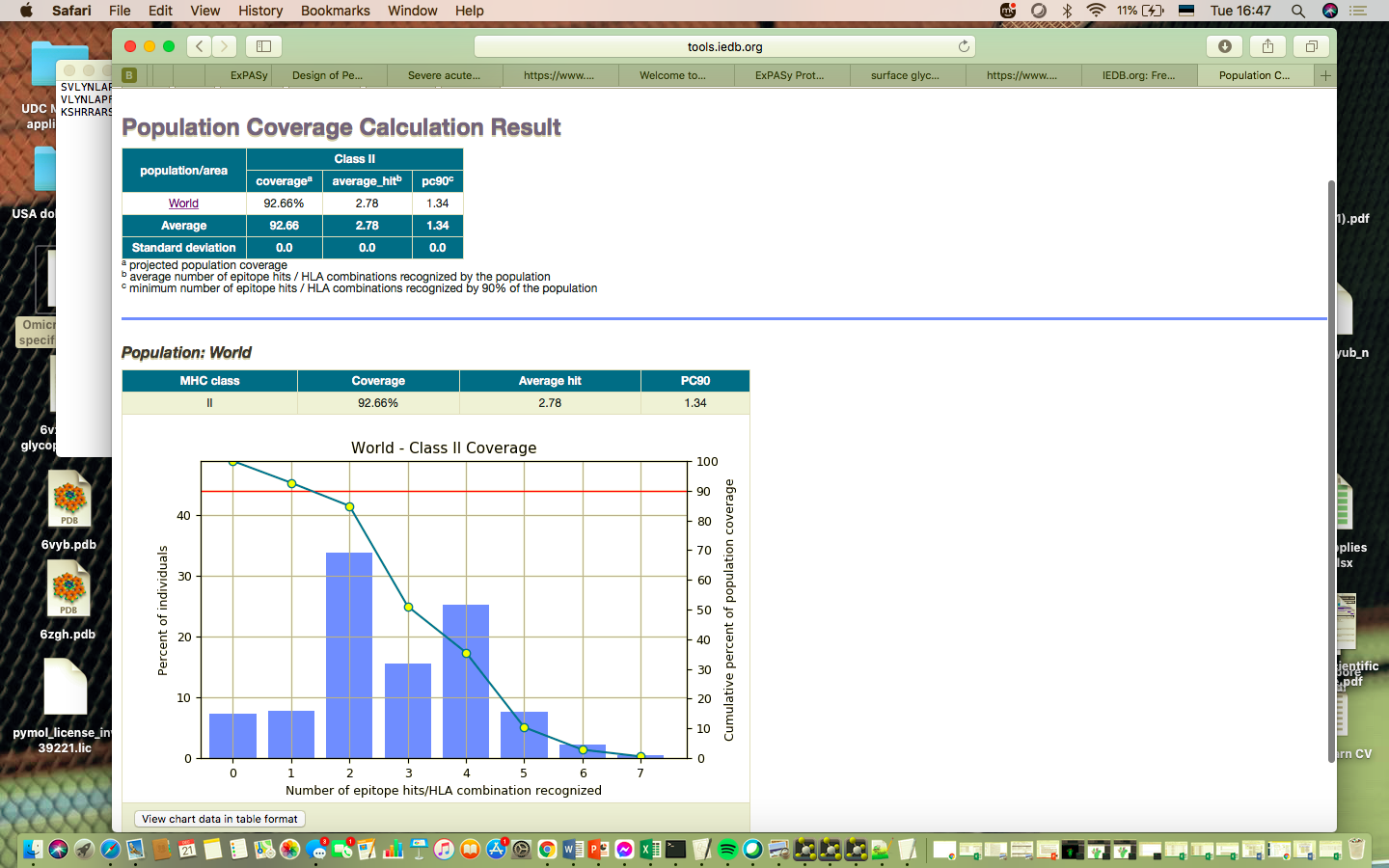

**Supplementary Figure S1.** World population coverage of top SARS-CoV-2 S protein T cell epitopes and their respective MHC Class I and II binding alleles. **(A)** World population coverage of top common CD8+ T cell epitopes across Omicron (BA.1 and BA.2) and other circulating SARS-CoV-2 variants including Alpha, Beta, Delta, Gamma, US variants (S protein mutations), and Cluster 5 mink variants. **(B)** World population coverage of top Omicron (BA.1) variant specific CD8+ T cell epitopes. **(C)** World population coverage of top common CD4+ T cell epitopes across Omicron (BA.1 and BA.2) and other circulating SARS-CoV-2 variants including Alpha, Beta, Delta, Gamma, US variants (S protein mutations), and Cluster 5 mink variants. **(D)** World population coverage of top Omicron (BA.1) variant specific CD4+ T cell epitopes. **(E)** World population coverage of Omicron (BA.1) variant specific CD4 peptides which overlap with CD8 peptides identified in this study.
